# From muscle fibres to muscle gears: How dynamic fascicle orientation and shape change impact skeletal muscle function

**DOI:** 10.1101/2025.06.15.659256

**Authors:** Matheus D. Pinto, James M. Wakeling, Javier A. Almonacid, Anthony J. Blazevich

## Abstract

Skeletal muscle architectural design strongly affects force-generating capacity and excursion range, and its functional importance to animal and human movement is well founded. Traditionally, this ‘structure−function relation’ has been inferred from architecture measurements with muscles at rest (from anatomical dissections or medical imaging techniques such as ultrasonography or magnetic resonance imaging) and muscle function tested under quasi-steady force conditions (e.g., isometric). A contemporary view recognises that, during active, dynamic contractions, muscles undergo load- and velocity-dependent changes in architecture and three-dimensional shape while remaining near-constant volume. Consequently, whole-muscle length change and velocity can become decoupled from fascicle length change and velocity, such that the whole muscle cannot be treated as a simple linear velocity transmission system but instead as a geared system where the input fascicle length change and velocity differ from the output whole muscle length change and velocity. This phenomenon is described by the concept of ‘muscle gearing’ and can be quantified as the ratio of the velocities or displacement of the muscle to the fascicle. Gear ratios are not fixed; instead, gear shifts with force and velocity, allowing the whole-muscle output to better match the mechanical demands of the task. Variable gearing is thought to emerge from load-dependent changes in fascicle angle and three-dimensional muscle deformations under volume-preserving and tissue constraints. Variable gearing has important functional implications, as it influences fascicle operating length and shortening velocity for a given task and thereby shifts where fascicles operate along their intrinsic force–length and force–velocity relationships. Through these effects, along with changes in the projection of fascicle force onto the muscle line of action as pennation varies, variable gearing can broaden whole-muscle force and power output across movement conditions. Accordingly, muscle gearing should be considered an integral component of a comprehensive framework for understanding dynamic muscle function in both normal and pathological states. This review aims to (i) provide a brief historical overview of how the muscle structure−function relation has been understood from resting muscle structure measurements and modelling and steady-force contractions to observations during dynamic, time-varying active contractions, and (ii) synthesise evidence on muscle gearing, including its measurement, functional consequences, and current understanding of the factors that influence it by reviewing findings from animal and human studies and raising hypotheses to explain the variation in gearing identified across muscle, species, and contraction conditions. Given the challenges in experimentally isolating the contribution of individual factors, we also employ a previously validated three-dimensional finite-element muscle model to interrogate the mechanical phenomena underpinning variable gearing and for the first time report effects from muscle activation. Finally, we briefly discuss the potential role of gearing in the observed changes in muscle function with ageing, injury, and exercise training. Overall, muscle gearing is a distinctive feature of skeletal muscles with important functional implications, and our modelling demonstrates that it is an emergent property of the physics of muscle contraction.

## I. Background

Skeletal muscles exhibit a wide range of contractile behaviours to meet mechanical demands across diverse movement tasks. A muscle can actively shorten to generate positive mechanical work during concentric contractions, contract isometrically to brace a body part or act as a strut, or actively lengthen to dissipate energy as a brake during eccentric (lengthening) contractions (Dickinson *et al*., 2000). This versatility is partly determined by the structural arrangement of fibres and fascicles because architecture determines both force-generating capacity and functional excursion range (Burkholder *et al*., 1994). Consequently, a muscle’s structure significantly influences its function, and this relationship has been a central focus in the study of animal and human movement for centuries (Lieber & Ward, 2011; Narici, Franchi & Maganaris, 2016).

Despite longstanding investigation in muscleLstructure function, much of our understanding of human muscle structure stems from observations made with the muscle at rest, typically obtained through anatomical dissections or imaging techniques (e.g., ultrasonography or magnetic resonance imaging), and from function tests performed under controlled isometric or during concentric (shortening) contractions. This is despite many years of research showing that muscles undergo dynamic changes in shape and structure during contraction to maintain near-constant volume and that many motor tasks require muscles to function eccentrically. Muscle shape changes occur across the structural scales, from the myofilament lattice space (myosin-actin filaments distance) (Williams *et al*., 2013) to fibre bundles (Smith, Fowler-Gerace & Lieber, 2011a) and whole muscle (Cobb, 2002) and directly influence muscle mechanical output. Thus, a comprehensive understanding of muscle function across contractile conditions must consider these deformations because the dynamic changes in muscle shape fundamentally impact muscle mechanical performance by directly decoupling fibre and fascicle velocity from whole muscle velocity in a process analogous to *muscle gearing* (Azizi, Brainerd & Roberts, 2008; Wakeling *et al*., 2011). For terminology clarity, here we (i) use fascicle reorientation to describe changes in fascicle angle during contraction, which are commonly referred to as fascicle (or fibre) rotation or pennation angle changes; (ii) use fascicle(s) because most methods described as ‘tracking fibres’ in practice track fascicles (or the perimysium surrounding fibre bundles) or marker-defined lines representing fibre-bundle motion; and (iii) avoid the term pennation angle when the muscle line of action or its force vector is unknown, such that the measured angle more precisely reflects fascicle angle rather than pennation *per se*.

The purpose of this review is to synthesise current understanding of muscle mechanics and function with a focus on muscle gearing (also termed belly or architectural gear ratio; see ref. Pinto *et al*., 2023 for methodological and nomenclature distinctions) and to outline its relevance to biological and health sciences. First, we summarise how the field has historically linked resting muscle architecture (structure) to contractile function. Second, we define ‘muscle gearing’ and explain how dynamic changes in fascicle orientation and muscle shape influence function under different contractile conditions. Third, we review proposed mechanisms underpinning muscle gearing and use a computational modelling framework to address limitations introduced by experimental design differences that can influence inferred gearing and complicate between-study comparisons. Within this framework, we present simulations using a previously published three-dimensional (3D) muscle model (Wakeling *et al*., 2020; Almonacid *et al*., 2024a) to interrogate the mechanical phenomena underlying muscle gearing. Fourth, we compare evidence from animal and human studies and discuss factors influencing muscle gearing *in vivo* that are not captured in isolated muscle preparations. Finally, we consider the potential contribution of muscle gearing to changes in contractile function with ageing, muscle and ligamentous injury, and exercise training.

## II. The muscle structure−function relationship: a brief historical overview

### (1) Importance of muscle architecture: the ‘classic’ view

Interest in skeletal muscle structure and function dates back to the Renaissance period (for detailed historical perspective, see refs. Narici *et al*., 2016; Kardel, 1990, 1994), when anatomical drawings already began to distinguish key architectural features. Classic textbooks and modern reviews typically classify muscle structure into one of two major categories: *parallel-fibred muscles*, with fibres oriented along the muscle’s line of action, and *pennate muscles*, with fibres oriented at an angle to it. These architectural differences have important functional implications because they impact contractile capacity, including force−length and force−velocity characteristics (Charles *et al*., 2022; Sacks & Roy, 1982). Muscle architecture is therefore widely used to infer a muscle’s force-generating capacity and functional excursion range (the distance over which a muscle can contract), and thus its mechanical output or performance (i.e., muscle function; Lieber & Ward, 2011; Lieber & Fridén, 2000).

In general, parallel-fibred muscles (with fibres aligned with the muscle’s line of action) have longer fibres than pennate muscles so they contain more sarcomeres in series, favouring larger excursion ranges and shortening velocity and a broader operating range along their force−length relationships (Lieber & Ward, 2011; Gans & Bock, 1965; Woittiez, Huijing & Rozendal, 1983). Parallel-fibred designs are thus thought to be useful for high-speed, dynamic movements (Blazevich & Sharp, 2005; Lieber & Fridén, 2000). Conversely, pennate muscles are thought to be better suited to higher force production because pennation allows more contractile tissue to be arranged in parallel within a given muscle volume (and tendon or aponeurosis attachment area), increasing physiological cross-sectional area and the number of sarcomeres in parallel (Blazevich & Sharp, 2005; Gans & Bock, 1965) and thereby conferring a mechanical advantage for force output (Rockenfeller *et al*., 2024). However, this architectural advantage comes with geometric and physiological costs; fibres run at an angle to the muscle’s line of action, decreasing the component of force in this line of action (Blazevich & Sharp, 2005; Gans & Bock, 1965; Zajac, 1992), and are shorter (relative to the muscle’s length) with fewer sarcomeres in series, which can limit excursion and shortening range relative to parallel fibred designs (Muhl, 1982; Lieber & Fridén, 2000; Alexander, 1983). Together, these principles underpin the classic structure−function and the trade-off between force capacity and shortening range and velocity.

#### (a) Limitations of the classical view

However, while useful, this ‘classical’ framework remains historically grounded on muscle architecture measured under fixed, non-contractile conditions (e.g., cadaveric dissections) and in geometrical (mathematical) models that relate resting structure to steady-state isometric conditions (e.g., *ex vivo* fixed-end preparations with clamps holding a muscle constant, or *in vivo* joint-level isometric tasks after force has reached steady state). In this framing, architecture is treated as largely invariant, and function is inferred without fully accounting for the transient reconfiguration of fascicles and whole-muscle shape during activation and dynamic contractions, nor for the dependence of these deformations on load and shortening velocity. Advances in force measurement and imaging (e.g., sonomicrometry, ultrasonography) now allow muscle architecture to be quantified during active contractions *in vivo* whilst recording time-varying forces (Blemker, 2023; Roberts & Dick, 2023), motivating a ‘contemporary view’ of muscle as a deformable system whose internal geometry changes during contraction. It is now recognised that these deformations are influenced by connective tissues and interactions with surrounding structures and pressures (Kelp *et al*., 2023; Eng, Azizi & Roberts, 2018; Roberts *et al*., 2019) and therefore muscle function cannot be accurately predicted solely from the architectural features of relaxed muscles.

### (2) Muscle shape change and fibre behaviour during active muscle contraction: a ‘contemporary’ view

#### (a) Early geometrical foundations and near-constant volume

A long-standing premise is that skeletal muscles behave as nearLconstantLvolume structures. As contractile elements shorten and force develops, the muscle expands transversely, and fascicle orientation and whole-muscle shape change. Early observations by Jan Swammerdam and Niels Stensen in the 1660s were instrumental in establishing this. Swammerdam demonstrated that touching the exposed nerve of an isolated frog gastrocnemius resulted in visible muscle swelling without apparent change in volume (retrieved from Kardel, 1996). Around the same time, Stensen (also known as Steno) developed a geometric framework for muscle contraction, representing pennate muscle using idealised parallelepiped models (reviewed in Kardel, 1994, 1990) (Figure 1A). Like Swammerdam, Stensen theorised that muscle volume remained unchanged during contraction (Figure 1B) and used geometry from Euclid’s *Elements* to show that shortening of obliquely oriented fibre is accompanied by transverse thickening while preserving volume. In doing so, he anticipated modern concepts of pennate muscle contraction mechanics. Together, Swammerdam’s empirical observations and Stensen’s geometric reasoning helped shift explanations of muscle contraction away from the prevailing recourse to animal spirits (Cobb, 2002).

**Figure 1.**
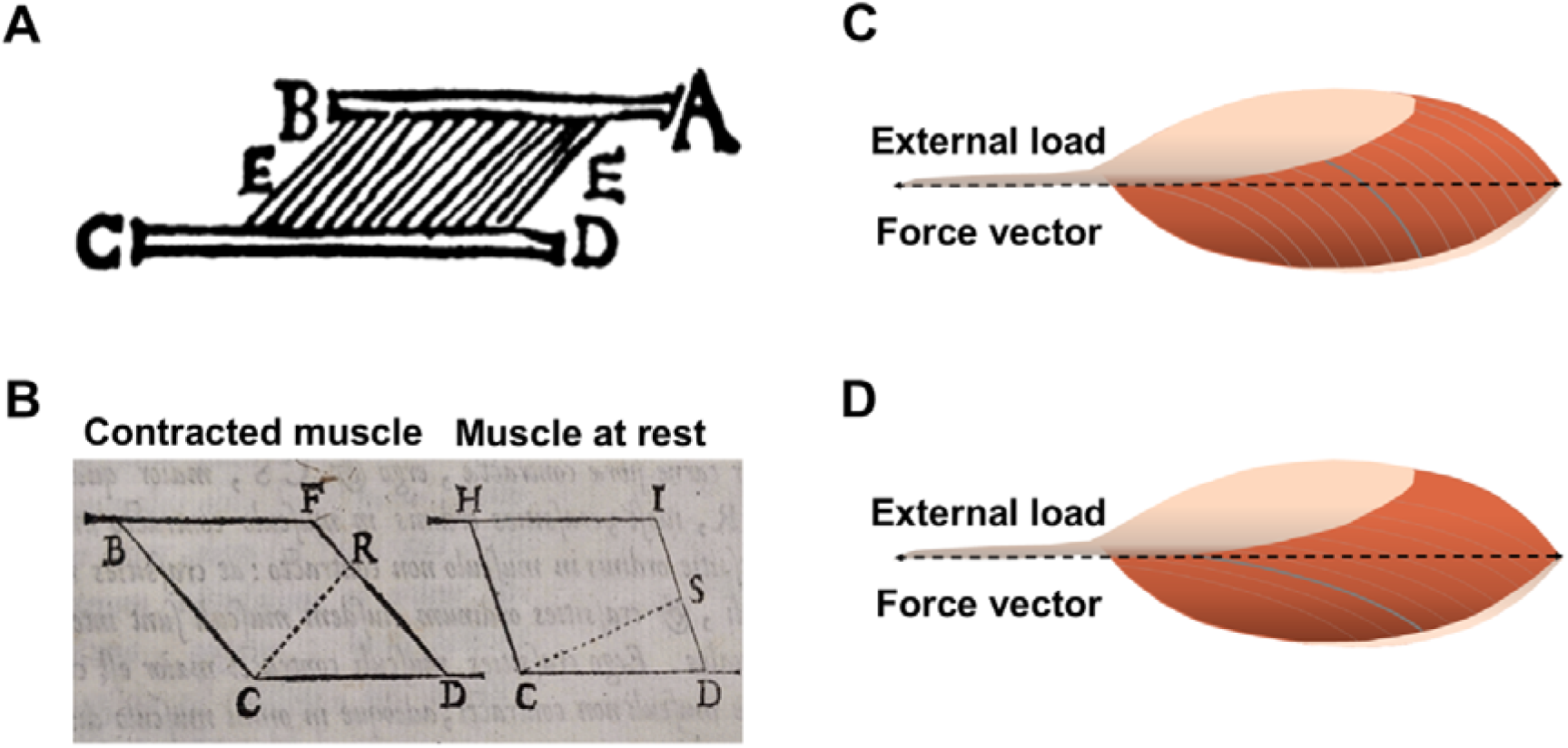
Historical origins and modern of pennate muscle geometry. (A) Geometric sketch by Niels Stensen (30 April 1663) proposed in correspondence letter with Thomas Bartholin. The sketch was informed by Stensen’s innovative practice of making longitudinal dissections along muscle fibres (rather than the more common transverse sections of his time), which revealed the internal organisation of obliquely oriented fibres and likely motivated his proposal that muscle contraction could be described using Euclidean geometry (Castel-Branco & Kardel, 2022). This idealised representation is now widely referred to as the unipennate actuator model and remains commonly used in biomechanical and physiological studies of skeletal muscle. Information and Figure retrieved from Kardel (1994; reprinted with permission). (B) Parallelogram representation from *Elementorum Myologiae Specimen* (1667) illustrating a muscle at rest and in the contracted state (adapted from Castel-Branco & Kardel, 2022; licensed under CC BY 4.0). Using Euclidean geometry, Stensen argued that the area muscle volume remains constant during contraction and that shortening reorientation of obliquely oriented fibres (from CS to CR) necessitates transverse thickening (muscle thickness) and a change in fibre obliquity, anticipating modern interpretations of pennate muscle bulging. Muscle thickness is defined here as the shortest (perpendicular) distance from point R to the lower aponeurosis (line CD), which increases during contraction without altering the area of the parallelogram. (C) Representation of a muscle with large and (D) small pennation angles (relative to the muscle’s line of action). Notice that, during muscle contraction, the change in segment length is expected to be greater than the fascicle length change in C, however there is a cost to peak force production because the component of fascicle force produced along the muscle’s line of action (external load/force vector) is reduced when compared to a muscle with lesser pennation, i.e., force along tendon = fascicle force × cos(pennation angle). This illustrates a fundamental trade-off between shortening capacity and transmission effectiveness.

#### (b) Early experimental hints of variable pennation angle and muscle bulging (1920s-60s)

Steno’s geometrical theories faced substantial resistance for over 300 years. In the 1920s, Beritoff (1925) demonstrated that, in pennate muscles, fibres shortened less than the whole muscle and its associated tendon because pennation angle increases during contraction. Beritoff (1925) further contended that the distance a whole muscle can shorten depends on concurrent thickening (bulging) of the muscle belly. These observations marked an early shift in understanding of muscle behaviour *during* contraction and were complemented by Feneis’ recording of fascicle angle changes during locomotion in transparent crabs ((Feneis (1935); reviewed in ref. Benninghoff & Rollhäuser, 1952)) and by Benninghoff & Rollhäuser (1952) experimental demonstrations that pennate architecture can increase whole muscle shortening distance, an idea later discussed in theoretical work of Gans & Bock (1965). However, experimental studies directly confirming and expanding Beritoff’s findings were not published in English until the 1980s.

#### (c) Observations and functional modelling of fascicle reorientation (1980s–1990s)

In 1982, Muhl exposed the rabbit digastric muscle and photographed fascicles and the whole muscle belly at rest and during active contractions at several lengths while recording tendon forces. Consistent with earlier predictions, Muhl (1982) observed changes in both fascicle length and angle during contraction, concluding that “for a given length of fibre, muscle excursion is enhanced by pinnation, relative to muscles in which the fibres lie in parallel to the line of muscle action.” Other experimental and modelling studies similarly showed that pennate muscles can exhibit greater whole-muscle than fascicle shortening distance and velocity and that this effect was not observed in parallelLfibred muscles of equal fibre length and physiological cross-sectional area (Huijing & Woittiez, 1983; Woittiez *et al*., 1984; Woittiez & Huijing, 1984). Modelling work attributed this dissociation largely to changes in fibre angle during shortening (Woittiez *et al*., 1984; Huijing & Woittiez, 1983; Otten, 1988). Despite these findings, many studies subsequently either neglected fascicle reorientation in estimates of force production (Bobbert, Huijing & van Ingen Schenau, 1986; Bobbert & van Ingen Schenau, 1990; Scott & Winter, 1991) or assumed fascicle angles remained fixed during contraction (Sacks & Roy, 1982; Spector *et al*., 1980; Scott & Winter, 1991).

The functional consequences of these assumptions were tested explicitly in the models developed by Scott & Winter (1991). They showed that if fibres shortened at a fixed pennation angle (i.e., no reorientation), fibre shortening velocity increased relative to muscle shortening velocity, *reducing* fibre force-generating capacity via the force–velocity relationship. In contrast, allowing pennation angle to increase during shortening reduced fibre shortening velocity relative to whole-muscle shortening, enabling *fibres* to operate more favourably on their force–length and force–velocity relations and allowing the *muscle* to function over a broader range of lengths and velocities (widening the *muscle’s* force−length and power−velocity relationships; as in Figures 2 and 3). Thus, pennation angle changes were predicted to decouple fibre and whole-muscle excursions and velocities, enhancing whole muscle excursion and shortening-velocity capacity (Scott & Winter, 1991). Similar conclusions were later supported experimentally in animal preparations (Zuurbier & Huijing, 1992, 1993).

**Figure 2.**
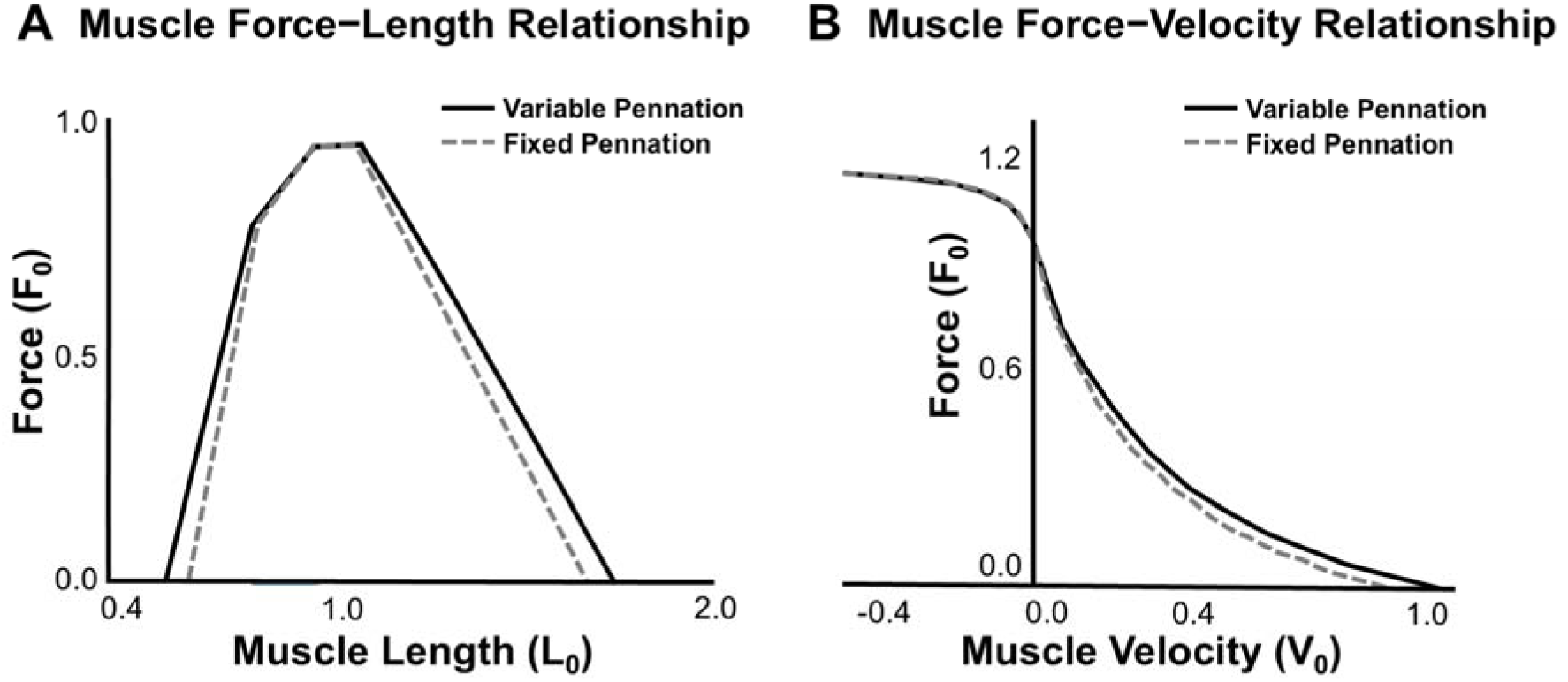
Effects of variable and fixed pennation (solid black and broken grey lines, respectively) on muscle (A) force−length and (B) force−velocity relationships. Figures adapted from Scott & Winter (1991). Reprinted with permission.

**Figure 3.**
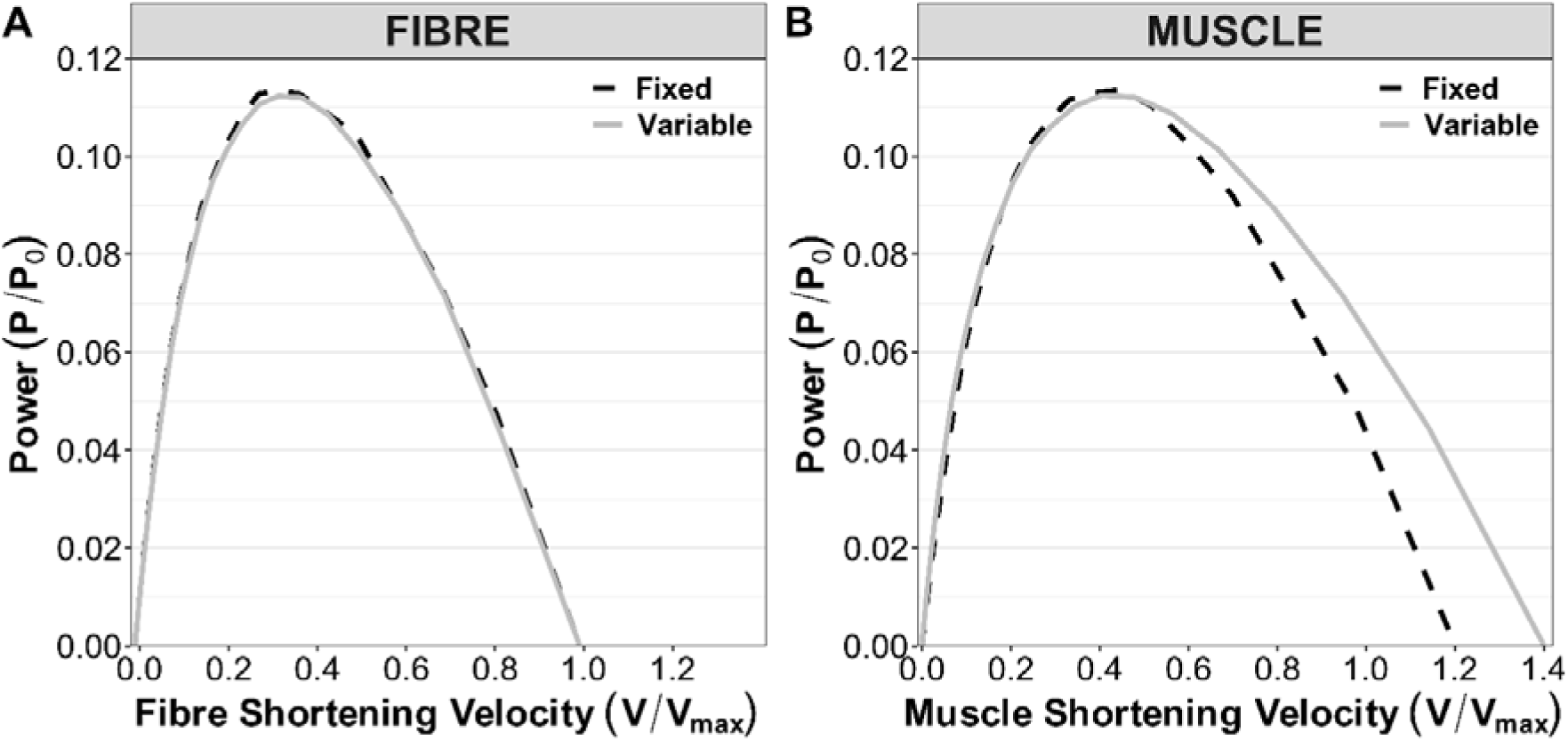
Effect of fixed (gear = 1.2; black dashed line) and variable (continuous grey line) gearing on fibre and muscle power−velocity relationships. Variable gearing broadens the range of muscle power production over a fibre velocity range. This figure is adapted from Fig. 6 in Azizi et al. (2008), where both force and shortening velocity are expressed relative to the maximal force and velocity, respectively. Power was calculated from the available shortening velocity and force data using values estimated using open available graph digitiser (WebPlotDigitiser, https://apps.automeris.io/wpd/).

#### (d) The missing piece: constraints on bulging in three dimensions

Although fascicle reorientation was progressively recognised as functionally important, the factors determining its magnitude remained unclear. Earlier work showed that pennate muscles thickened during shortening and thinned during lengthening, and modelling showed that neglecting thickness change substantially biased muscle and tendon length changes for a given fascicle shortening (Huijing & Woittiez, 1983; Woittiez *et al*., 1984). Yet many models treated thickness changes as small and therefore neglected the mechanical work required for a muscle to expand against surrounding tissues (e.g., aponeurosis, skin, adjacent muscles and bones) during shortening (Reinhardt *et al*., 2016; Wakeling *et al*., 2020). Alexander’s theoretical framework suggested that angled collagen fibres in fish myosepta could constrain lateral body movement and bulging during shortening (Alexander, 1969); however, he and others (Otten, 1988; Gans & Bock, 1965) often assumed that near-constant volume constraints (Baskin & Paolini, 1967) implied a constant distance between the tendon plates (aponeuroses) and thus similar transverse deformations in thickness and width. Collectively, studies from the 1920s (Beritoff, 1925) through the 1980’s provided evidence that muscles could bulge in thickness during contraction and this could influence fascicle reorientation and muscle and tendon excursions, and that surrounding non-contractile tissues likely shaped these deformations. What remained unresolved was whether, and how, connective tissues modulated deformations across the two transverse axes, and if this affected fascicle□muscle decoupling in ways that meaningfully influenced muscle function.

#### (e) Connective tissue constraints, 3D deformation, and architectural gearing

Using salamander segmented musculature and modelling, Azizi, Gillis & Brainerd (2002) demonstrated that the decoupling between muscle segment and fascicle strains depended on whether shortening was accompanied by bulging in thickness (direction normal to the superficial and deep aponeuroses; z-direction) or width (the orthogonal transverse dimension; y-direction). In their model, segment strain matched fascicle strain when thickness decreased, exceeded fascicle strain when thickness was held constant, and was greatest when thickness increased. They proposed that connective tissues could ‘steer’ bulging between orthogonal axes to preserve constant volume (Azizi *et al*., 2002); consistent with this, morphological analyses revealed that myoseptal collagen fibres were angled to help resist deformation in the width direction (mediolateral; y-axis), thereby prompting bulging in the thickness direction (dorsoventral; z-axis). This ‘bulge control hypothesis’ provided a structural basis for greater muscle segment strain relative to fibre strain (Azizi *et al*., 2002).

Subsequently, Brainerd & Azizi (2005) proposed that the decoupling between fascicle and segment strain could be characterised by the architectural gear ratio (AGR; segment strain/fascicle strain) and showed in simulations that AGR increases, and muscle segment shortens more for a given fibre shortening, with higher initial fascicle angle with bulging conditions that favour thickness expansion due to larger increases in fascicle angle. Azizi *et al*. (2008) then demonstrated that AGR varied with the mechanical demands of the task, with higher AGR occurring in low-force, high-velocity contractions to enhance muscle shortening velocity relative to fascicles, whereas lower AGR occurring in high-force, low-velocity contractions to improve force output. Together, these findings provided a novel perspective that dynamic architectural changes and whole-muscle shape change are functional determinants of muscle performance, not only minor geometric details.

## III. Muscle gearing: definition and functional importance

Muscle gearing refers to a mechanism that affects muscle length and velocity during contraction through fascicle reorientation and associated three-dimensional muscle shape change (Azizi *et al*., 2008; Wakeling *et al*., 2011). As fascicle orientation (angle) changes during contraction, fascicle velocity becomes partially decoupled from whole muscle (or muscle segment) velocity (Azizi *et al*., 2008; Brainerd & Azizi, 2005), and this gearing effect can be characterised by the ratio of whole muscle to fascicle length change or velocity (e.g., Δ*L*_muscle_/Δ*L*_fascicle_ or *V*_muscle_/*V*_fascicle_; see ref. Pinto *et al*. (2023) for nomenclatures and methods). Gear is >1.0 when muscle velocity exceeds fascicle velocity, is 1.0 when the muscle shortens or lengthens at the same velocity as its fascicles and is <1.0 when fascicles shorten or lengthen faster than the whole muscle. In pennate muscles, the muscle gear ratio is generally ≥1.0 (Eng *et al*., 2018), although this has not been systematically described and is a focus of the current review (see Table 2).

Changes in muscle shape during contraction significantly affect the magnitude of fascicle reorientation and therefore the magnitude of fascicle and whole muscle velocity decoupling. Because muscle shape changes and reorientation are governed by the force exerted by a muscle and the load imposed on it (Eng *et al*., 2018; Azizi *et al*., 2008) the extent of this decoupling changes in response to the mechanical demands of the contraction. Therefore, the gear in which a muscle operates depends on the force produced during contraction (Azizi *et al*., 2008; Eng *et al*., 2018). Muscles have been shown to operate in high gear when contracting against light loads or when generating lower force and possibly higher velocity (Azizi *et al*., 2008). These contraction conditions are characterised by substantial changes in fascicle angle and bulging in muscle thickness, resulting in large muscle length changes (Azizi *et al*., 2008). Conversely, muscles shift to low gear when contracting against heavier loads and producing high forces with lower velocity (Azizi *et al*., 2008). Under these conditions, fascicles reorient less and muscle shape changes more predominantly in width, resulting in less muscle length change relative to fascicles (Azizi *et al*., 2008) since fascicles are more aligned to the tendon (Figures 1C-D). Whilst it remains unclear as to whether gearing is indeed affected by force or rather velocity (Wakeling *et al*., 2011; Azizi *et al*., 2008; Dick & Wakeling, 2017), changes in gear, i.e., *variable gearing*, allow muscle force output to better match the demands of a task by shifting fibre behaviours such that, like in other geared systems, gear ratio will adjust to favour force production under high loads and movement speed under low loads (Eng *et al*., 2018; Roberts *et al*., 2019; Dick & Wakeling, 2017).

LoadLdependent fascicle behaviour and gearing have important functional implications. During *concentric* contractions, variable gearing broadens the muscle’s functional range (Figure 3), shifting and perhaps more importantly broadening (widening) the power−velocity relation (although it has no effect on the peak power). Whilst shifts in gear may allow muscles and their fibres and fascicles to partially overcome the force−velocity constraints associated with cross-bridge kinetics (time-dependent cross-bridge detachment and re-attachment; Rome *et al*., 1999; Podolsky, Nolan & Zaveler, 1969) by enabling them to operate more optimally within their force−length and force−velocity relations, most of the benefit of variable gearing appears to manifest at the whole muscle level, as depicted in Figure 3. A *muscle* that maintains constant gear in response to variations in load (and thus muscle force) displays a narrower power−velocity relation that is largely dictated by and closely mirrors the *fibre* and *fascicle* power−velocity relations. In contrast, a muscle that appropriately changes gear displays a broader whole□muscle power−velocity relation that extends beyond the limits of the fibres and fascicles (e.g., Figures 3 and 11). During *eccentric* contractions, variable gearing allows muscles to lengthen more than the fascicles at any given external load and is therefore a feature of pennate muscles that reduces active fascicle strains and strain rates that potentially minimises muscle damage and the risk of strainLrelated injury during eccentric (lengthening) contractions (Pinto *et al*., 2023; Azizi & Roberts, 2014). Given that variable gearing allows the muscle to broaden its performance range, understanding the mechanisms of gearing may have a range of clinical and health applications.

However, despite the functional importance of variable gearing, our understanding of the mechanisms and structures influencing fascicle reorientation, three-dimensional muscle shape changes, and gear shifts remains an evolving area of research. A range of studies have investigated different aspects of the mechanisms of variable gearing, including the role of pennation, threeLdimensional expansion, tissue properties, intramuscular pressure, external pressure and external work on the muscle surface. In the following section, we present evidence for the inferred and deduced mechanisms for these different aspects.

## IV. Mechanisms of muscle gearing

Initial evidence supporting variable gearing predominantly comes from studies of isolated muscle preparations, where supramaximal and short-lived stimulations are delivered to muscles to evoke supramaximal and non-transient activations, in addition to bioinspired muscles. These studies overall suggest that this feature is not imposed by the nervous system (Azizi *et al*., 2008; Azizi & Roberts, 2013; Sleboda, Roberts & Azizi, 2024; Wang *et al*., 2022). Instead, a growing body of evidence indicates that variable gearing and muscle shape changes during contraction result from the incompressible nature of skeletal muscle tissues and their composite structure (Eng *et al*., 2018). Variable gearing appears to be mediated by how fascicle (vectorial) forces, elastic connective tissue elements, and other structural constraints (e.g., fat tissue) affect, and therefore interact with, one another (Roberts *et al*., 2019; Eng *et al*., 2018). These passive (non-contractile) structures are found at the levels of the fibre, fascicle, and whole muscle level and are tightly packed within a volume constrained structure (the whole muscle belly); their arrangement, in addition to their quantity, appears to contribute to gearing. Fascicle orientation, i.e., pennation angle, is also an important factor influencing muscle gearing and, as such, it influences load-dependent gear change (Azizi *et al*., 2008; Brainerd & Azizi, 2005). Experimental and theoretical evidence for the influence of each of these factors on the dynamics of muscle gearing is discussed below.

### (1) Pennation angle

Resting pennation angle strongly influences muscle gear partly because muscles with a higher pennation at (rest and) the start of muscle contraction display greater fascicle reorientation. This greater change in angle more effectively contributes to muscle length change for a given amount of fascicle shortening (Eng *et al*., 2018; Brainerd & Azizi, 2005) (as illustrated in condition A, Figure 4, Table 1). However, high gear should also be observed in muscles with greater initial pennation, even when undergoing *equal* fascicle reorientation compared to those with less initial pennation. This is because a given change in fascicle angle results in a larger change in muscle (segment) length when initial pennation is higher. Specifically, the end points of a given fascicle would move more in the *longitudinal direction* when initial pennation is high but more *orthogonally* (in the thickness direction) when pennation is lower. This behaviour can be explained by the geometrical principles of sine and cosine laws. For a given change in angle, there will be more significant increases in the height (i.e. muscle thickness) and less reduction in the adjacent (muscle) segment length when initial angle is low (although this will vary according to fascicle length and its changes). Conversely, when initial angle is high, the same change in angle will contribute more to the reduction of the segment length and less to height increase. As muscles often increase in thickness during dynamic muscle contraction, this is an expected behaviour, although in some cases muscles may also exhibit little to no change in thickness during contraction *in vivo* (Maganaris, Baltzopoulos & Sargeant, 1998; Narici *et al*., 1996; Wakeling *et al*., 2011). However, this behaviour should occur even if muscle thickness remains constant during contraction, as muscles with greater starting pennation will work in higher gear. Thus, the initial pennation angle is an important factor influencing gear (Azizi *et al*., 2008; Wang *et al*., 2022; Brainerd & Azizi, 2005).

**Figure 4.**
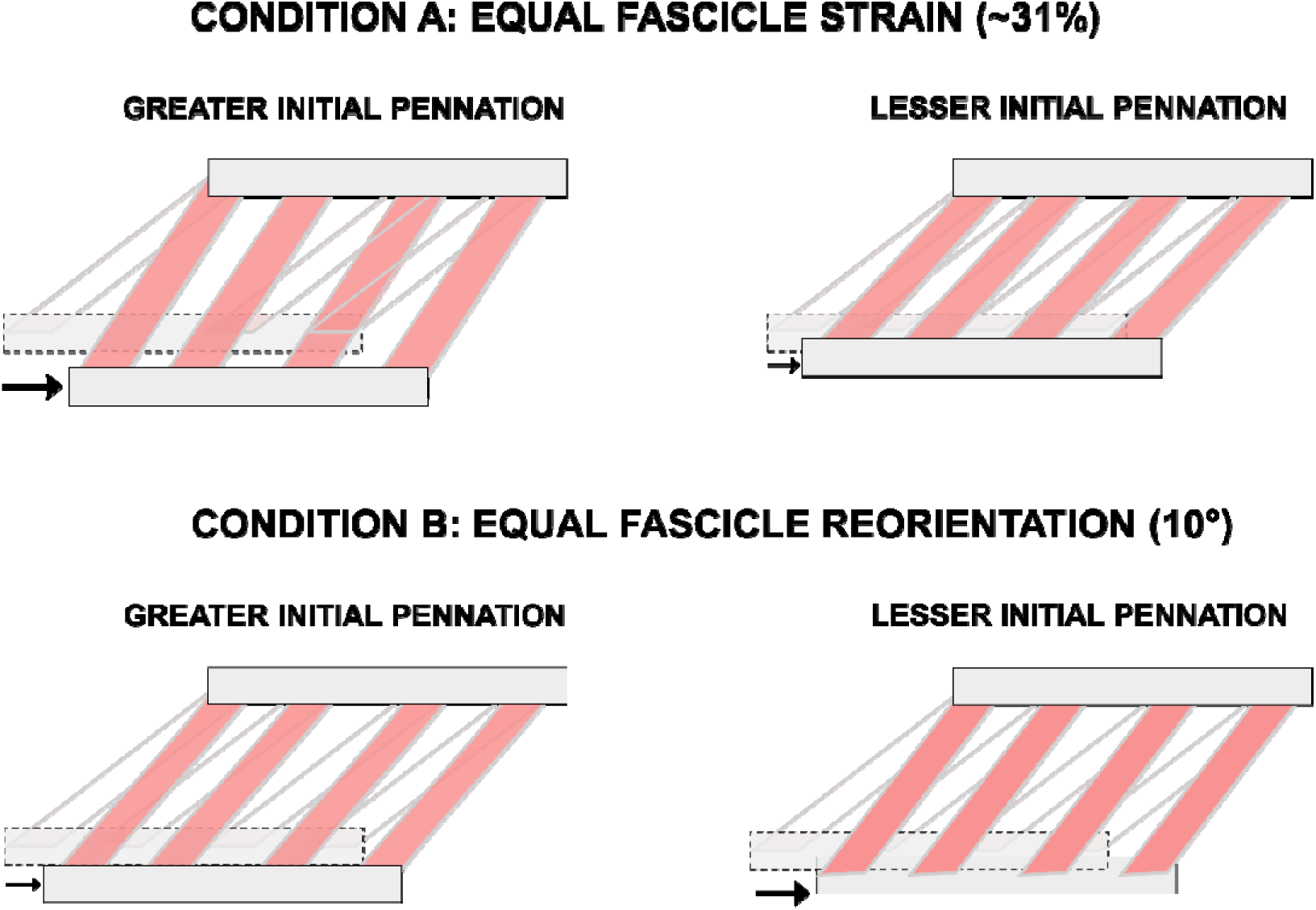
Schematic of a muscle block (a section of the muscle belly) illustrating the effect of initial pennation angle on changes in pennation angle, muscle segment length, and muscle gear during contraction with equal fascicle shortening (condition A) and equal fascicle reorientation (condition B). The architectural characteristics of each example are given in Table 1. Black arrows indicate the magnitude of muscle segment shortening. Note: figures are not drawn to scale; they are representations of the phenomena.

**Table 1.** Architectural parameters for the biplanar sketch in conditions A and B in Figure 4.

CONDITION A - Equal fascicle shortening, 20% thickness increase
|  | Higher initial pennation |  |  |  |  | Lower initial pennation |  |  |  |  |
| --- | --- | --- | --- | --- | --- | --- | --- | --- | --- | --- |
| | $L_f$ | $L_t$ | $\beta$ | $L_{seg}$ | Gear ratio | $L_f$ | $L_t$ | $\beta$ | $L_{seg}$ | Gear ratio |
| Before shortening | 6.55 | 3.40 | 31.3 | 5.60 |  | 6.55 | 2.16 | 19.3 | 6.18 |  |
| After shortening | 4.50 | 4.10 | 64.6 | 1.94 | **1.80** | 4.50 | 2.60 | 35.1 | 3.70 | **1.22** |
| Change ($\Delta$) | 2.03 | 0.68 | 33.3 | 3.66 | | 2.03 | 0.43 | 15.8 | 2.48 | |

CONDITION B - Equal fascicle reorientation, 20% thickness increase
|  | Higher initial pennation |  |  |  |  | Lower initial pennation |  |  |  |  |
| --- | --- | --- | --- | --- | --- | --- | --- | --- | --- | --- |
| | $L_f$ | $L_t$ | $\beta$ | $L_{seg}$ | Gear ratio | $L_f$ | $L_t$ | $\beta$ | $L_{seg}$ | Gear ratio |
| Before shortening | 6.55 | 3.40 | 31.3 | 5.60 |  | 6.55 | 2.16 | 19.3 | 6.18 |  |
| After shortening | 6.19 | 4.10 | 41.3 | 4.65 | **2.61** | 4.41 | 2.16 | 29.3 | 4.63 | **1.25** |
| Change ($\Delta$) | 0.36 | 0.68 | 10.0 | 0.95 | | 1.24 | 0.43 | 10.0 | 1.55 | |

$L_f$ = fascicle length (cm); $L_t$ = muscle thickness (cm); $\beta$ = pennation angle (degrees); $L_{seg}$ = muscle segment length (cm); Change ( $\Delta$ ) = absolute difference between parameter before and after shortening.

Despite evidence indicating that higher pennation is associated with greater reorientation and larger gear ratios (Tijs, Konow & Biewener, 2021; Brainerd & Azizi, 2005), it is important to note that muscles with higher pennation do not always exhibit a higher gear (Azizi & Brainerd, 2007). This inconsistency illustrates muscle gearing as a multifactorial phenomenon (as further discussed below).

### (2) Fascicle forces driving muscle bulging: balance between radial expansion-driven fascicle reorientation and thickness compression

When pennate muscles shorten to generate active force, fascicles undergo transverse deformations to maintain near-constant volume (Baskin & Paolini, 1967). As fascicles are packed within a volume-constrained muscle space, running adjacent to each other and with tight packing density, the radial expansion of adjacent fascicles pushes neighbouring fascicles away from each other, and this occurs as fascicles reorient to greater angles (Dick & Wakeling, 2017; Figure 5, blue arrows). Decreased radial expansion of the fascicles concurs with reduced fascicle reorientation (Ryan *et al*., 2019). Similarly, decreasing the spacing between neighbouring McKibben actuators reduces actuators reorientation and eliminates variable gearing likely due to decreased transverse forces acting on adjacent actuators (Sleboda *et al*., 2024).

**Figure 5.**
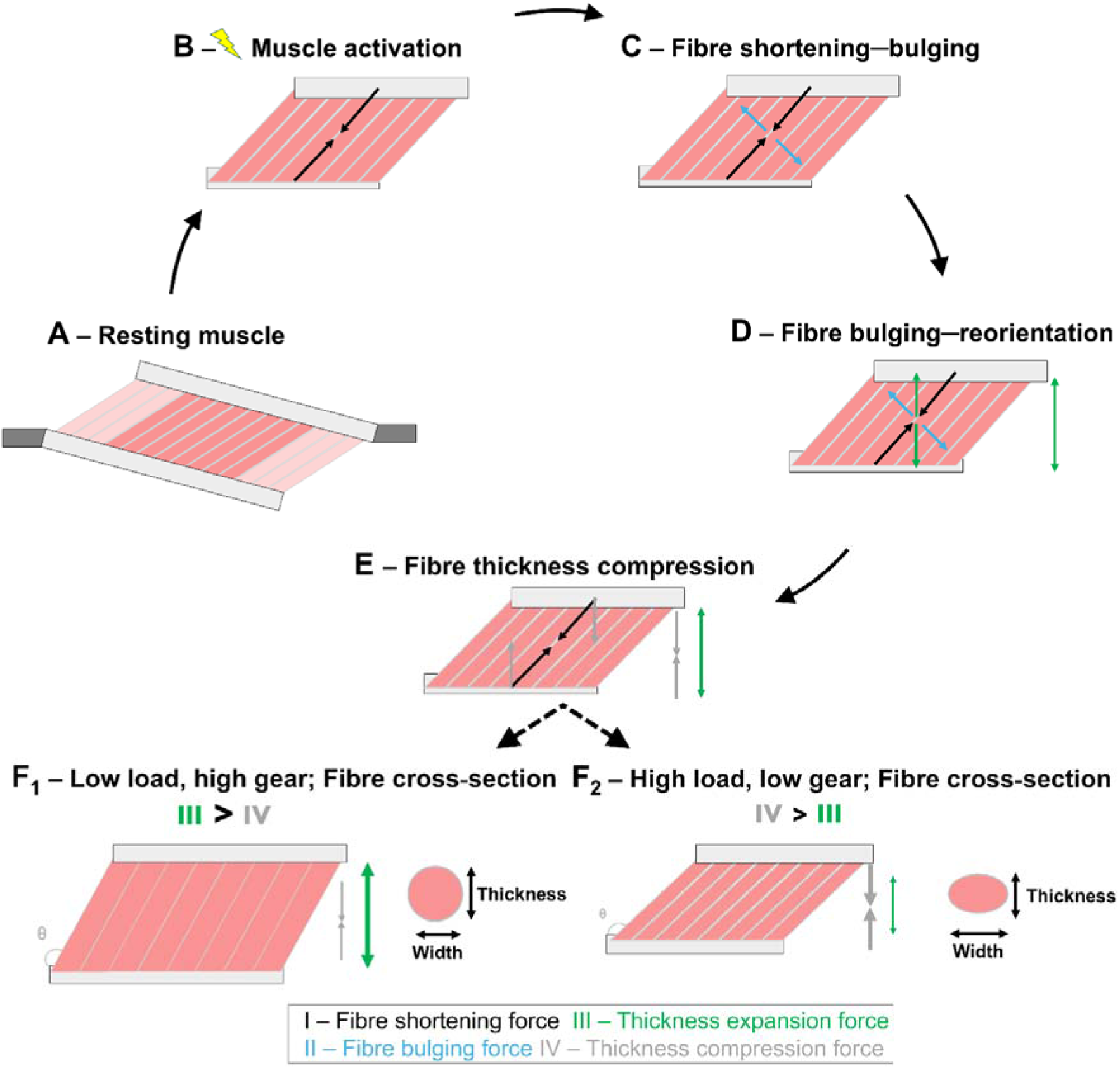
Architectural, muscle shape, and gear changes during muscle contraction. (A) Illustration of pennate muscle architecture in a non-activated, relaxed state, with fibres packed within a volume-constrained muscle belly. In panels B to F, a localised portion of the muscle is re-drawn as an idealised muscle “block” (the non-shaded element bounded by aponeurotic sheets) to simplify the geometry and visualise the relevant force components. (B) Upon activation and contraction, the contractile components within the fascicles generate forces that shorten the fascicles along their line of action (black arrows). (C) Since fascicles remain nearly constant volume, fascicle shortening is accompanied by radial expansion, which produces forces orthogonal to the fascicles’ line of action (blue arrows). Orthogonal fascicle forces push neighbouring fascicles away and pennation increases i.e., fascicles rotate. (D) Increases in fascicle angle creates a force that separates the aponeuroses apart, increasing the muscle belly thickness (green arrow). (E) This force is counteracted by a component of the fascicle shortening force that acts to reduce muscle thickness and bring the aponeuroses closer together illustrated in grey. The change in muscle thickness is partly mediated by the balance between fascicle bulging forces that promote thickness expansion and orthogonal fascicle shortening forces that promote thickness compression. This balance is dictated by the amount of force fascicles need to produce in response to the external load. (F) When the load is low, the thickness forces are smaller than the forces imposed by fascicle reorientation, resulting in greater changes in fascicle angle and higher muscle velocity relative to fascicle velocity, as illustrated in (low load, high gear). (G) When the external load is high, the fascicles generate greater force, compressing the muscle in thickness and resulting in less changes in fascicle angle and slower, more similar whole muscle velocity to that of the fascicles. The muscle operates at low gear in this scenario.

Changes in fascicle angle creates a force that acts to separate the aponeuroses and increases their distance in the thickness direction (Figure 5, green double headed arrow). This force is simultaneously counteracted by the component of the fascicle shortening force that acts to reduce muscle thickness and bring the aponeuroses closer together (thickness compression; refer to Figure 5, grey arrows). The change in overall muscle belly thickness is therefore influenced by the balance of these two forces. A visual representation of this process is provided in Figure 5A□C.

Variable gearing occurs because fascicle radial expansion and the fascicle thickness compressive force are determined by the force fascicles produce in response to the external load (Eng *et al*., 2018; Roberts *et al*., 2019; Azizi *et al*., 2008). When the load is low and fascicles generate low forces and fascicle radial expansion occurs approximately equally in both thickness and width (assuming isotropy). This equally distributed radial expansion promotes increases in fascicle angle, with this reorientation being facilitated because the fascicle thickness compressive force is smaller than the force produced by fascicle radial expansion that induces fascicle orientation and increases thickness (Figure 5D). However, under high loads in which fascicles generate greater force, muscle radial expansion is constrained in the thickness direction because the component of fascicle shortening force pulling the aponeuroses together is stronger, limiting thickness expansion but not resisting fascicle and belly width-wise expansion that occurs to maintain a constant volume, as in Figure 5E. With less opportunity for belly thickness expansion and thus fascicle reorientation, muscle length change is closer to the fascicle length change, and the muscle gear is low (∼1.0; Figure 5F_1_). Importantly, observations that asymmetrical transverse deformations of fascicles (Randhawa & Wakeling, 2018; Kinugasa *et al*., 2012) and muscles (Hodson-Tole & Lai, 2019) occur have been documented human muscles *in vivo*.

### (3) Connective tissue material properties and architecture

The extent and direction of muscle shape change are not solely determined by the balance of fascicle forces. Instead, the extracellular matrix (ECM) provides multiscale constraints that can limit bulging, bias deformations toward width or thickness, and thereby modulate fascicle reorientation and muscle gearing (Eng & Roberts, 2018; Azizi *et al*., 2017; Azizi *et al*., 2002; Roberts *et al*., 2019; Holt *et al*., 2016; Alexander, 1969).

#### (a) Endomysium and perimysium: resistance to bulging and shear-mediate sliding

The endomysium forms an organised, honeycomb-like collagen network that envelops and links adjacent muscle fibres (Purslow, 2020). It comprises longitudinally oriented tubular “shells” aligned to fibre axes (Sleboda, Stover & Roberts, 2020; Gillies & Lieber, 2011), with collagen arranged in a wavy, quasi-random pattern (Purslow & Trotter, 1994; Rowe, 1974). The perimysium links these structures and surrounds fibre bundles in a mesh-like sheath (Gillies & Lieber, 2011; Purslow, 2020) that includes helically wound collagen cables (Sleboda *et al*., 2020; Gillies & Lieber, 2011) crossing within a multilayer, crimped pattern (Gillies & Lieber, 2011). These structural differences have been proposed to support distinct mechanical roles (Purslow, 2020), with shear-compliant perimysium potentially facilitating inter-fascicle sliding and transverse fascicle expansion but the shear-stiff endomysium serving to maintain tight fibre alignment and facilitate force transfer. Collagen fibre length and orientation change as muscle fibres and fascicles shorten (Purslow, 1989; Trotter & Purslow, 1992). Consistent with a functional role for constraining bulging, fibre-wound cylinder models show that collagen angle and area influence the coupling between radial expansion and longitudinal strain (Azizi *et al*., 2017), as illustrated in Fig. 6. Accordingly, endomysial and perimysial architecture is expected to affect bulging and interfascicular shear, with downstream effects on fascicle reorientation and muscle gear.

**Figure 6.**
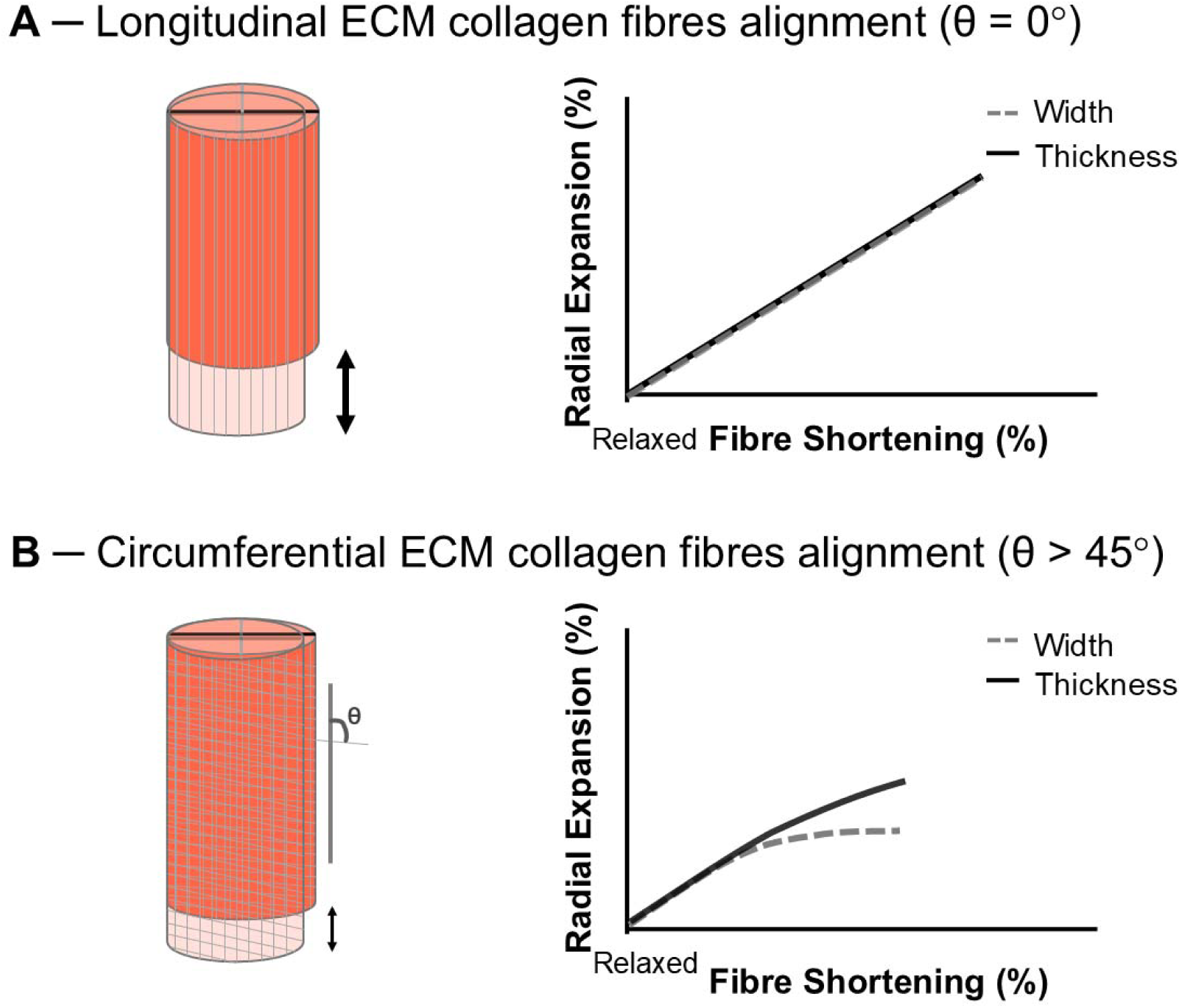
Hypothetical example illustrating the effect of extracellular matrix (ECM) collagen fibre orientation on fibre or fascicle strain and shape changes in width (grey) and thickness (black), assuming constant fibre or fascicle volume. Relaxed fibres (light pink) are overlaid on active fibres (orange) to illustrate deformation and longitudinal fibre strain (double-headed arrow). In panel A, the ECM collagen fibres align with the fibre’s longitudinal axis at a negligible angle (θ = 0°), primarily resisting longitudinal fibre shortening (Azizi et al., 2017). If the ECM behaves isotropically, providing uniform resistance in all directions, width and thickness strains remain equal, preserving circular cross-section. However, ECM anisotropy may lead to slight deviation from perfect circularity. In panel B, ECM collagen fibres are more circumferentially orientated at a high pitch angle (θ > 45°), restricting both longitudinal fibre strain and radial expansion. In this condition, ECM anisotropy is assumed, so width and thickness strains may differ and result in asymmetric deformation. If circumferential constraints (high hoop stiffness) preferentially limit width expansion, then width changes will be less than thickness changes, altering the fibre’s cross-sectional shape. The grey line represents width strain, and the black line represents thickness strain.

#### (b) Epimysium and aponeurosis: whole muscle constraint that redirect bulging

The epimysium and aponeurosis form robust collagenous sheets that envelop the muscle and provide the insertion surface for pennate fascicles (Sleboda *et al*., 2020; Rowe, 1974). They resist tensile stress effectively but are less suited to resist compressive loads (Morrow *et al*., 2010; Purslow, 2020), such as fascicle forces that compress the muscle belly in thickness. During contractions in which the muscle is compressed in thickness and expands in width, these structures are stretched to resist tensile stress (Morrow *et al*., 2010; Eng *et al*., 2018). This *resistance to width expansion* is thought to *promote thickness expansion* to maintain constant muscle volume, thereby influencing muscle shape change during contraction (Eng *et al*., 2018; Eng & Roberts, 2018).

#### (c) Evidence linking ECM constraints to muscle gearing

Holt *et al*. (2016) reported that older rats operated in near-constant (intermediate) gear across forces, whereas young rats shifted from high to low gear as force increased; older rats also exhibited greater transverse aponeurosis stiffness and greater longitudinal stiffness of fibre bundles. Higher gear during high-force contractions was attributed to increased resistance to thickness compression due to stiffer aponeurosis, which reduced thickness compression and width expansion. Consequently, older rats exhibited larger increases in fascicle angle and operated at a higher gear despite the high force. In adult birds, small longitudinal incisions in the aponeurotic sheet, presumably reducing transverse stiffness and allowing width-wise bulging, reduced gear during high-force contractions but did not abolish variable gearing (Eng & Roberts, 2018). The lower gear at high forces was hypothesised to result from increased thickness compression and width expansion, indicating an inability to resist deformation and to prevent muscle flattening during contraction. Together, these observations align with finite element models showing that changes in aponeurosis stiffness affect fibre geometry and behaviour (Rahemi, Nigam & Wakeling, 2014), and support the view that ECM constraints can influence muscle shape changes and load-dependent muscle gear.

### (4) Mechanical pressure: internal and external sources

Muscles generate internal pressure during contraction (Hill, 1948), which can influence muscle performance by modifying stress distribution and resisting sarcomere shortening forces (Daggfeldt, 2006). This internal pressure has been empirically measured and modelled (Van Leeuwen & Spoor, 1993; Sejersted *et al*., 1984; Heukelom, van der Stelt & Diegenbach, 1979), and is thought to arise from a combination of forces within the muscle, including those produced by fibre bulging and internal fluids as well as structural mechanical interactions between closely packed fibres and fascicles. Intramuscular pressure is further modulated by external factors such as the forces exerted by adjacent muscles (Reinhardt *et al*., 2016), bones, intermuscular fat, and the ECM, which either passively resist transverse expansion or, in the case of adjacent muscles contracting to generate lateral pressures, actively compress the muscle. External compressions or loads provide further resistance to fibres and fascicles bulging and muscle shape changes (Ross *et al*., 2020; Wakeling, Jackman & Namburete, 2013; Siebert *et al*., 2016). Here, we consider internal and external sources of pressure separately because they arise from different structures, yet both appear to impose transverse constraints on contracting muscle and are therefore plausible modulators of muscle shape change and gearing.

#### (a) Internal: the potential role of fluid and structural pressures

Skeletal muscles contain a substantial amount of fluid within muscle fibres, interstitial space, and blood vessels surrounding them (Bhave & Neilson, 2011), and they can influence internal pressure and fascicle dynamics in complex ways. While fluid pressure is often cited as a factor influencing intramuscular mechanics, its precise role in muscle shape changes and force transmission remains debated. Some interpretations consider muscles as a hydrostatic system, where fluid movement is responsible for redistributing stress and mechanically interacts with elastic elements surrounding muscles to modulate muscle shape changes during contraction (Eng *et al*., 2018; Kier, 2012). An alternative view is that muscles behave and can be modelled as a poroelastic (biphasic) material (Wheatley *et al*., 2017, 2018), where internal pressures (solid-fluid interactions) contribute to structural deformation without requiring significant fluid displacement.

During contraction, fibres and fascicles shorten and expand in girth (Figure 5), which is thought to generate internal pressures that influence stress distribution. If fibres are implicitly considered as fluid-filled hydrostats, then the hydraulic forces would redistribute contractile stress between longitudinal and radial directions (Eng *et al*., 2018). Hydraulic stresses have been suggested to redirect compressive fascicle forces to promote transverse expansion that stretches the surrounding connective tissues. Connective tissues resist this stretch, resulting in fluid redirection (i.e., hydraulic stress redistribution) that counters thickness compression (Eng *et al*., 2018). This dynamic interplay between fascicle forces, connective tissues, and fluid has been suggested to involve flow of energy across the fascicular and whole-muscle levels and influence their behaviours (Roberts *et al*., 2019; Eng *et al*., 2018). However, the flow of energy linking longitudinal to transverse deformations can also be explained by simple structural models (Otten, 1988; Siebert, Till & Blickhan, 2014a) and more complex continuum models of muscle in which fluid is absent (Wakeling *et al*., 2020; Ryan *et al*., 2020a; Blemker, Pinsky & Delp, 2005). Poroelastic models, which account for both solid and fluid interactions (Wheatley *et al*., 2018; Böl *et al*., 2022), may be an additional framework to understand these energy dynamics.

Experimental evidence nevertheless suggests that fluid-induced pressures can influence muscle mechanics. Increasing muscle fluid volume (via hypotonic solutions) increases resistance to stretch (Sleboda, Wold & Roberts, 2019; Sleboda & Roberts, 2017) and can generate passive shortening forces even in the absence of electrical input (Wold, Sleboda & Roberts, 2021), potentially via pressure-induced changes in ECM collagen orientation that resist longitudinal stretch (Sleboda *et al*., 2019; Sleboda & Roberts, 2017) and facilitate muscle shortening (Wold *et al*., 2021). This behaviour resembles that of ‘hydrostatic skeletons’, like worms, where internal pressures modulate stiffness and connective tissue structural arrangement to influence movement mechanics (Kier, 2012). Variable gearing in McKibben pneumatic actuator arrays provides a further analogue of pressure-driven transverse expansion and reorientation (Azizi & Roberts, 2013; Sleboda *et al*., 2024; Wang *et al*., 2022). Pressurised elastic tubes within braided sleeves expand radially and shorten as sleeve fibres reorient, increasing array thickness and promoting transverse bulging and load-dependent gearing. However, these devices shorten by increasing volume and therefore do not replicate the near-constant-volume constraint of skeletal muscle (Baskin & Paolini, 1967).

Further research is needed to determine how fluid, or its absence as in dehydrated muscle preparations, modulates muscle shape change and variable gearing. Dehydration can induce cell stiffening and shape deformation (Guo *et al*., 2017), raising the question of whether such effects reflect altered fluid-mediated pressures or changes in solid-phase material behaviour. Anisotropic deformations can, in principle, emerge from internal stress redistribution and energy transfer within structured solids, even without bulk fluid movement. Poroelastic models offer an emerging framework for capturing this (Wheatley *et al*., 2017, 2018) and be used to understand variable gearing, but current evidence suggests that ECM material properties and structural constraints likely play a more dominant role in muscle shape changes and gearing than bulk fluid movement, particularly in complex muscle groups that experience mechanical interactions and compressions as they contract (see below).

#### (b) External pressure: surrounding structures and mechanical loading

Structures external to the muscle (e.g., neighbouring muscles, intermuscular fat, and bones) can constrain transverse expansion. Evidence supporting this effect comes from the reductions in the rate of force development and maximal force from the application of transverse external compression to the surface of the contracting rat medial gastrocnemius (Siebert, Günther & Blickhan, 2012; Siebert *et al*., 2014b; Siebert *et al*., 2016; Siebert *et al*., 2014a). This likely resulted from forces acting at the muscle’s surface needing to be counteracted by the redirection of muscle work to the surface, resulting in reduced internal force generation capacity along the line of muscle action (Wakeling *et al*., 2020; Siebert *et al*., 2014a). In humans, external compression via elastic bandage also reduces belly gear and attenuates changes in fascicle angle and muscle thickness during contraction (Wakeling *et al*., 2013). Across compression paradigms, adding external load alters muscle geometry (Ryan *et al*., 2019; Stutzig *et al*., 2019; Monte, 2021), reduces force production (Ryan *et al*., 2019; Stutzig *et al*., 2019; Siebert *et al*., 2018), and increases energetic cost (Monte & Zamparo, 2023), with stronger effects seen when compression is applied along the muscle thickness axis than the width axis (Ryan *et al*., 2020b). Empirical data and finite element modelling indicates that compression-related force reduction is greater at shorter muscle lengths (Sleboda & Roberts, 2020) or when resting pennation angle is low (Ryan *et al*., 2020a).

In muscles operating within their muscle package *in vivo*, intermuscular pressures of 10–70 kPa have been measured at the surface of contracting muscles (Reinhardt *et al*., 2016; Bernabei, van Dieën & Maas, 2017). Synergists may partially compensate for such constraints via coordinated shape changes. During concentric plantar flexion contractions, the medial gastrocnemius can decrease in thickness while lateral gastrocnemius increases in thickness (Randhawa, Jackman & Wakeling, 2013), a pattern replicated across tasks and cohorts (Randhawa & Wakeling, 2018; Wakeling *et al*., 2011; Kelp *et al*., 2021; Kelp *et al*., 2023). This coordinated behaviour suggests that architectural differences between synergists may redistribute deformation such that as one muscle thins the other ‘fills the space’, potentially stabilising compartment shape and moderating pressure-related effects on function.

Finally, muscle size and mass may influence deformation dynamics via inertial effects. Simulations indicate that internal mass can reduce muscle shortening velocity, particularly when activation is submaximal (Ross & Wakeling, 2021, 2016). Together, these findings demonstrate that externally applied boundary pressures (e.g., compressive garments and added external mass and loads) can constrain transverse deformation and thereby modulate muscle shape change and load-dependent gearing independent of internal fluid mechanisms. However, the extent to which intermuscular pressures and inertial effects from muscle mass contribute to variable gearing and muscle shape changes require further investigation.

## V. Computational modelling of muscle gearing – an emergent property of pennate muscles

The evidence reviewed above supports that variable gearing is a multifactorial phenomenon that appears to result from the complex non-linear interactions of (i) pennation angle, (ii) contractile element vectorial forces, (iii) connective tissue architectural and material properties, and (iv) internal and external pressures. Given this complexity, we introduce a theoretical framework that allows these effects to be considered in synergy. Such an approach allows interrogation of knowledge gaps that have not yet been experimentally mapped, and to address more general mechanistic questions about gearing.

To understand the fundamental mechanisms underlying muscle gearing, it is ideal to quantify the independent influence of each factor reviewed herein. Alternatively, one could remove some or all structural or biomechanical features to determine whether gearing still occurs. These approaches are challenging to implement experimentally; however, computational (*in silico*) modelling offers a framework for understanding such complexities and examining the specific factors underlying muscle gearing. It enables ‘what-if’ experiments (DiSalvo & Blemker, 2024), allowing specific factors to be manipulated independently or instead explored using a simplified base model to reduce computational time. This approach also facilitates testing unresolved questions, such as whether dynamic changes in muscle shape and gearing are influenced by force, velocity, activation, or a combination of these factors. Unlike animal studies, which typically involve supra□maximal muscle activation, computational models allow exploration of a range of activation conditions that reflect *in vivo* contractions.

Therefore, to probe deeper into the mechanisms underlying muscle gearing and changes in muscle shape during contraction, we ran a series of dynamic (rather than quasi-static) simulations using a 3D finite element muscle model (Wakeling *et al*., 2020; Ryan *et al*., 2020a; Almonacid *et al*., 2024a; Almonacid *et al*., 2024b). Unlike previous quasi-static models, this model explicitly accounts for the internal effects of muscle mass and fibre shortening velocity. The model geometry was styled from the average (young) rat medial gastrocnemius muscle reported by Holt *et al*. (2016), resulting in a block of muscle tissue of 18.6 × 6.58 × 6.95 mm with an initial pennation angle of 26.1° and a mass of 0.9 g. For each simulation we stretched the block to 110% of its initial (resting) length then ramped the activation to a predetermined level and finally shortened the block in an isovelocity manner while it was still active. When the activated block shortened back to its initial length (L_0_), the mean muscle velocity, mean fibre velocity, fibre pennation angle, and the muscle block thickness and width through its midsection were measured. The gear ratio was computed at this timepoint, and the force at the end of the muscle was determined in the line-of-action of the muscle block. Figure 7 displays the simulation time-series and the muscle block and fibre-level visualisations at key events of the protocol.

**Figure 7.**
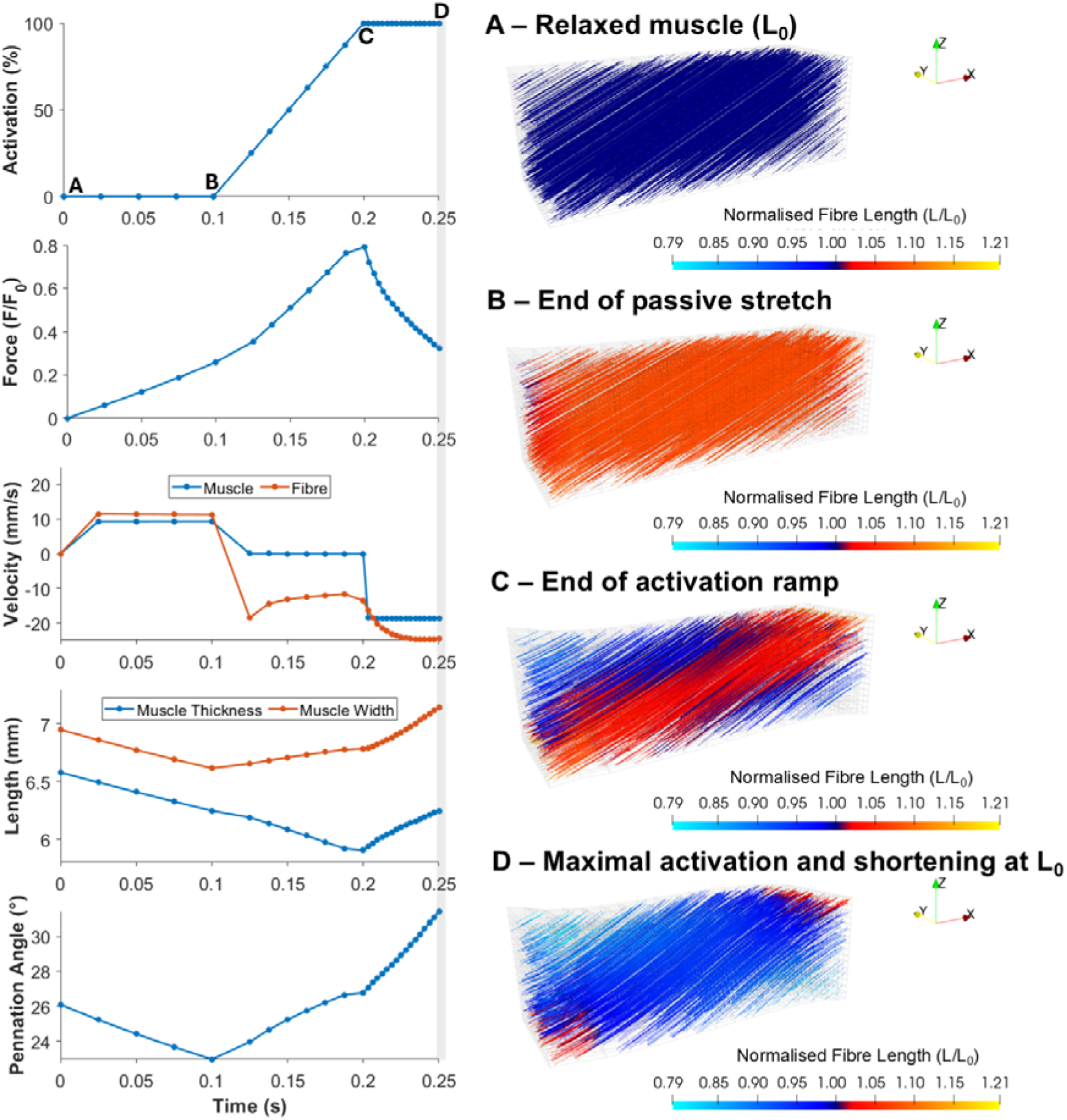
Dynamic simulation of muscle gearing during active shortening. Left panel: Time□series plot showing the sequence of events from resting (A) to, passive stretch (B), activation ramp whilst isometric (C), and active shortening at constant activation (D) for activation (% of maximum), muscle force (normalised to optimal length), fibre and muscle velocities, muscle shape (muscle thickness and width at the midsection), and fibres pennation angle. The grey region denotes the time in which variables were measured for analyses reported below. The muscle block was first passively stretched to 110% of its initial length (L_0_), followed by a ramped activation phase and then actively shortened back to L_0_ under constant velocity. Right panel: Visualisations of the muscle block shape and fibres displacement distribution (coloured-coded) at the four time points illustrating both fibre and muscle shape changes during the key events. Arrows in each panel indicate the orientation of the coordinate system (x: anterior□posterior/ muscle block length; y: medial□lateral/ muscle block width; z: superior□inferior/ muscle block thickness).

We formulated the muscle model in the same manner as previously described (Wakeling *et al*., 2020; Almonacid *et al*., 2024a), as a fibre-reinforced composite biomaterial. The one-dimensional fibres encoded the contractile elements of the muscle and had active and passive forceLlength and forceLvelocity properties typical of a Hill-type muscle model (with a maximum unloaded strain rate of 5 s^-1^, and a maximum isometric stress of 200 kPa). These fibres were encased in a three-dimensional volume representing the tissue properties of the muscle; the tissue properties were a homogenised representation of the base material that included both cellular and extra-cellular (ECM) components. The contribution of intramuscular fat was not included in the homogenised base material, and so these simulations inform us of the contribution of fat-free muscle tissue to variable gearing. Tissue properties included the energetic cost of the muscle changing shape, small volume changes (<1%), and the kinetic energy of the tissue. The internal pressure within the muscle was additionally incorporated into the equations system (Almonacid *et al*., 2024a) for solving the gearing. Simulations were run with a time interval between steps such that 16 time-steps of simulation were computed during active shortening (i.e., after the muscle reached 110% of its initial length and the predetermined activation level). At each time step, the total strain energy of the system was minimised (i.e. the energetically optimal solution was found for that time).

When the model was run, the fibres shortened and also rotated to increase in pennation as the muscle block shortened (Fig. 7). The rate of fibre reorientation increased with both muscle shortening velocity and increased activation (Fig. 8). The fibre shortening velocity increased with muscle velocity but decreased at higher activations (Fig. 8). The muscle gear (mean muscle block velocity/mean fibre velocity) decreased with muscle force (Figs. 9 and 10), increased with muscle velocity (Figs. 9 and 10), and increased with activation (Fig. 9). There was a velocity-dependence to muscle gearing that was independent of force (Fig. 9). This effect is consistent with previous reports on human muscle (Wakeling *et al*., 2011), and emerged because the simulations tested a range of activation levels that have not been previously possible in *in-situ* or *in-vitro* animal preparations.

**Figure 8.**
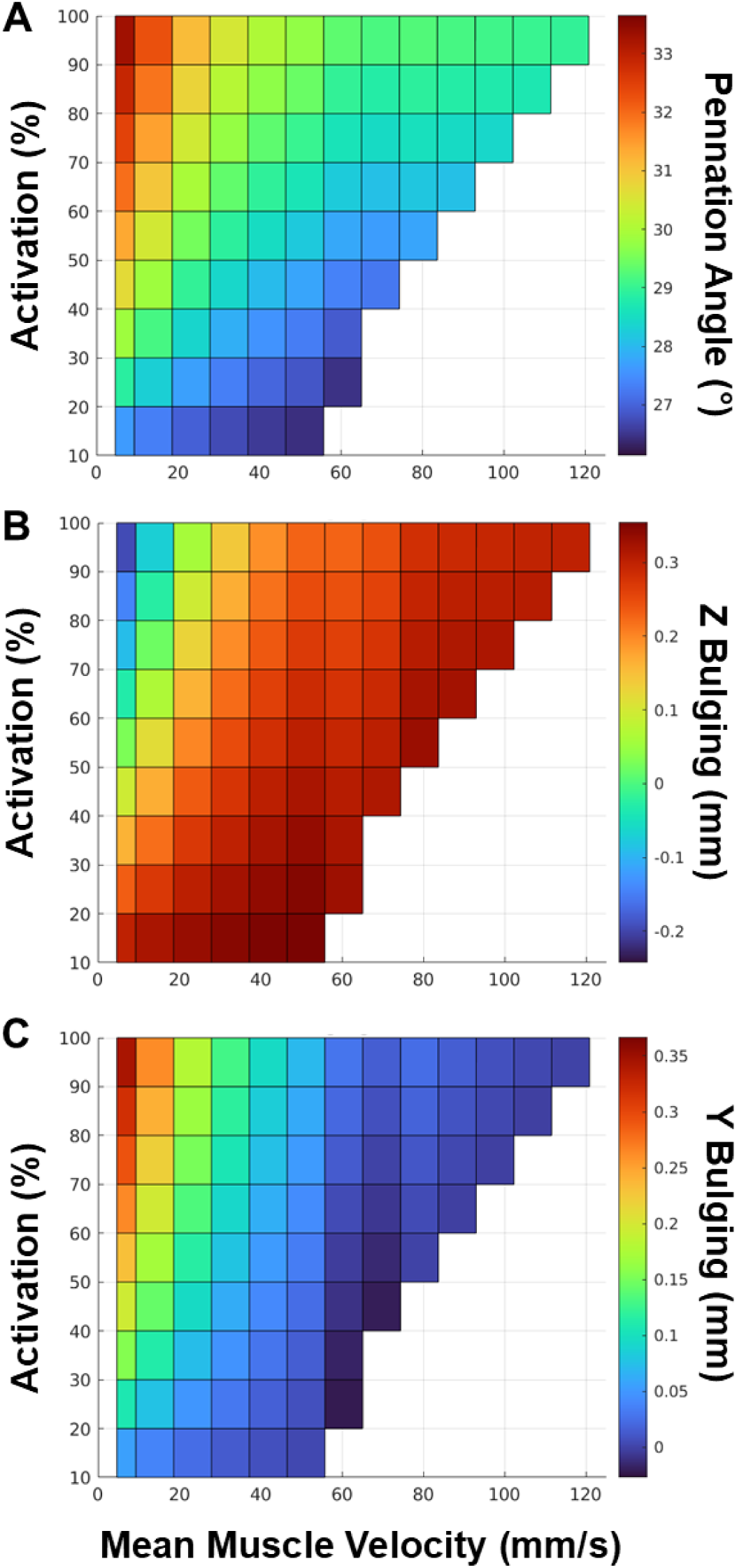
Activation- and velocity-dependent changes in muscle pennation angle and transverse shape during active shortening. (A) Pennation angle increases with greater activations and lower shortening velocity. (B) Muscle bulging along the thickness direction (Z□axis, orthogonal to the line of action) decreases with activation, suggesting compression at high activation levels and expansion at low activation levels, with a small influence of velocity. (C) Muscle bulging along the width direction (Y□axis, orthogonal to thickness) increases with both activation and shortening velocity. The schematic below/above illustrates the modelled muscle block before and after shortening, depicting both architectural and dimensional changes corresponding to panels A-C.

**Figure 9.**
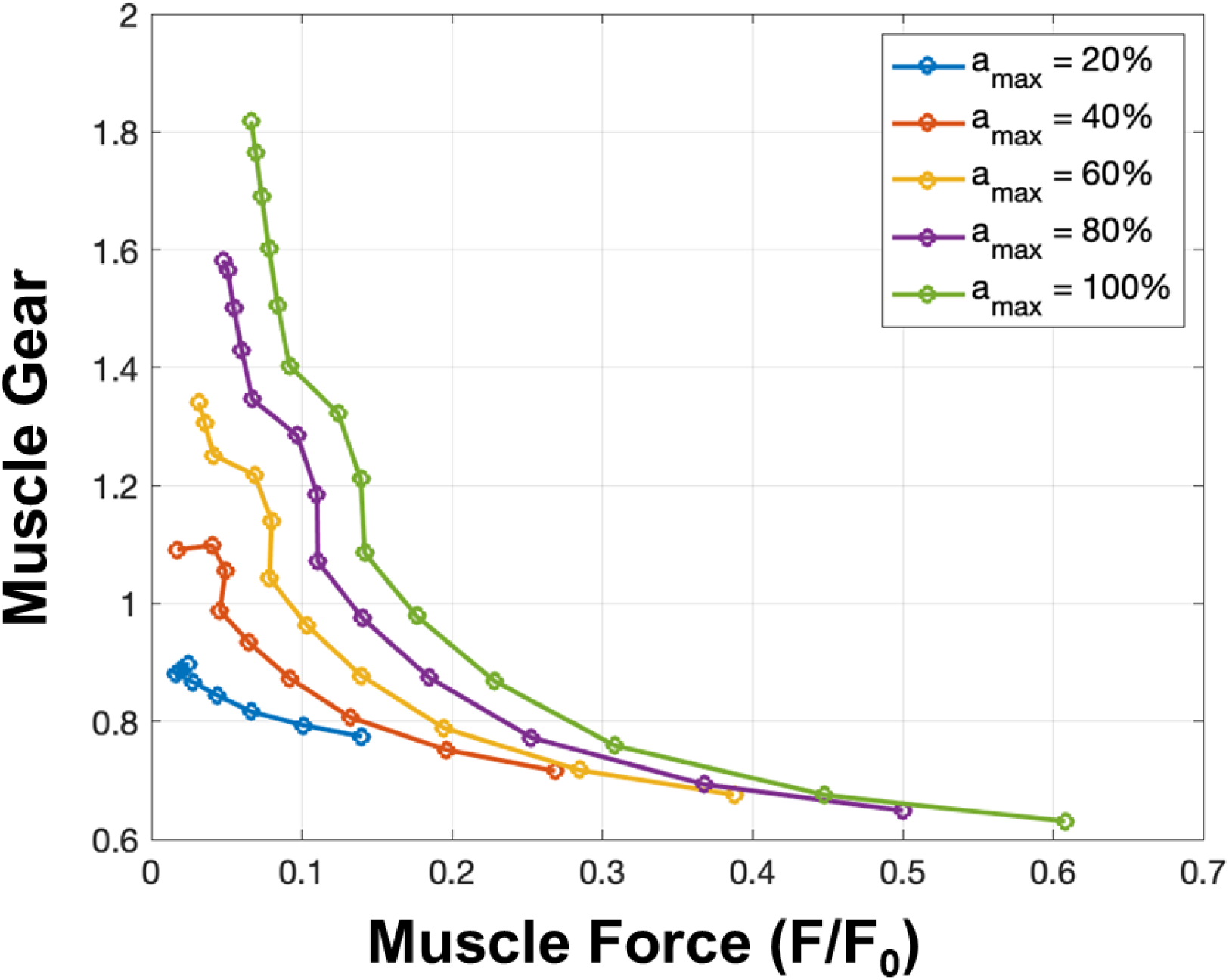
Effect of activation level on the muscle gear−force relationship. Muscle gear ratio decreases non-linearly with increasing normalised muscle force (F/F0), with higher levels of maximal activation (a□□□) enabling higher gear at low force outputs. This dynamic interaction between activation and the mechanical gearing of muscle fibres during force generation suggests that muscle gear is activation-dependent.

**Figure 10.**
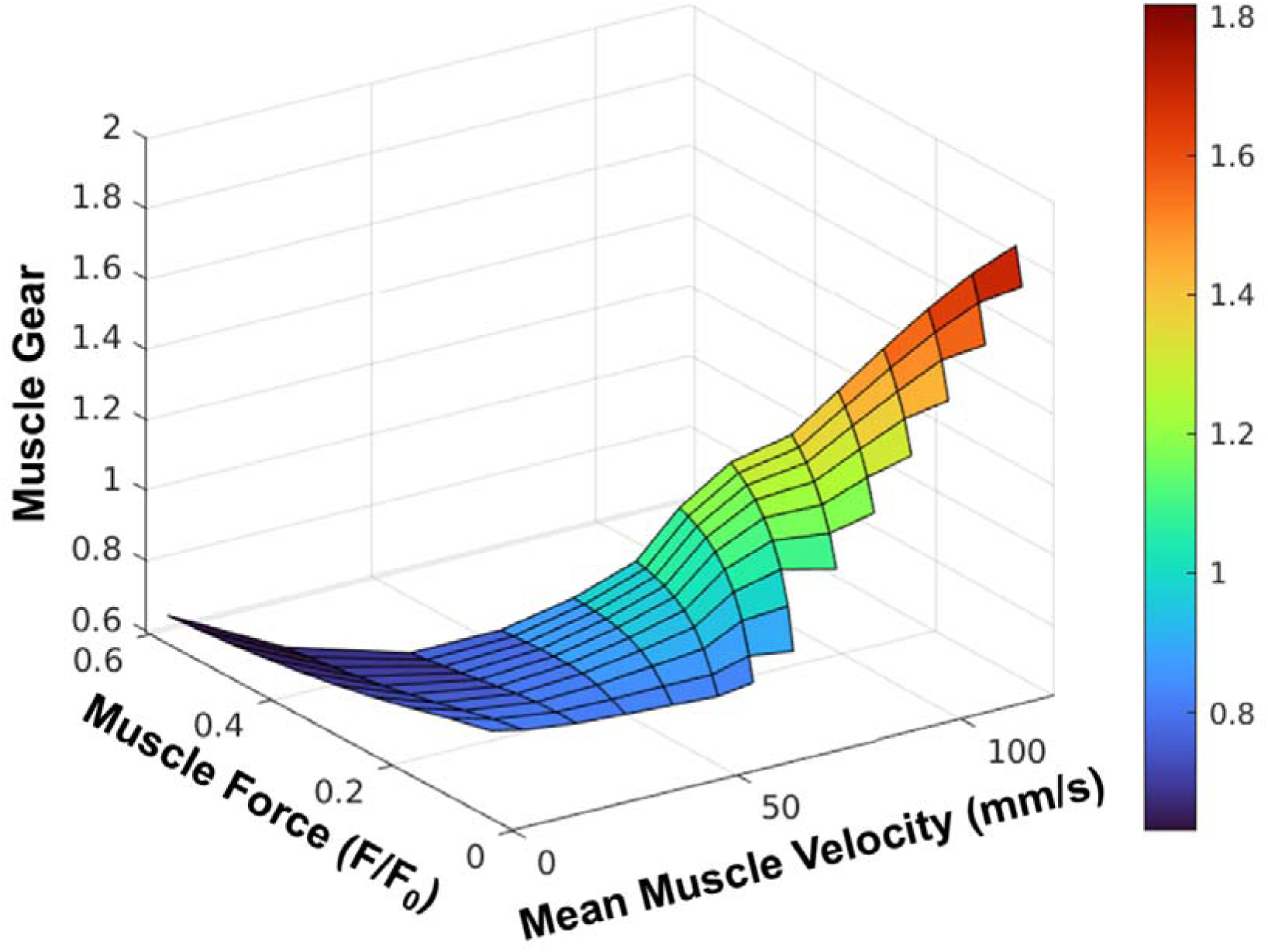
Relationship between muscle gear, muscle force, and mean muscle velocity during active muscle shortening. Muscle gear ratio decreases with increasing force but increases with mean fibre velocity, largely independent of force level. Gear values are particularly high during high shortening velocities, when gearing would be beneficial to limit fascicle length changes (i.e., both velocity and length operations) during rapid contractions.

As the muscle block shortened, it bulged in its midsection in an attempt to conserve its volume. At low velocity and high activation, the thickness of the muscle (z-direction; in the fibre xz-plane) decreased a small amount, but at higher velocities the thickness increased (Fig. 7). The width of the muscle block was maintained or increased in all conditions (Figure 7). Thus, the highest belly gearing occurred with a large increase in thickness but minimal change in width, whereas the lowest gearing coincided with a decrease in thickness by an increase in width. These findings are consistent with the model proposed by Azizi *et al*. (2008).

Of note from these simulations is that all the general features of muscle gearing emerge from the physics of the contractile elements and the muscle tissue properties alone, and do not need effects from external tissues to occur. Importantly, additional effects from tissues such as aponeurosis (Rahemi *et al*., 2014; Rahemi, Nigam & Wakeling, 2015; Wakeling *et al*., 2020), ECM (Konno, Nigam & Wakeling, 2021a) and intramuscular fat (Rahemi *et al*., 2015) can certainly be incorporated into this modelling framework but were not necessary to address the gearing mechanisms described herein. Further, the simulations predict similar gearing effects that have been reported for maximally activated muscles in *in situ* animal studies but also now predict the more nuanced effects that occur at submaximal activations, as is the case during human movement. This simulation framework shows how force, velocity, and activation independently affect muscle gear. The exact state of the tissue when the measures were probed depends on the energy balance at each condition, which in turn depends on the complex interaction between tissue stiffness, compressibility, and mass as well as the activation state and fibre contractile properties.

## VI. Variable gearing *in vivo*: what have we learnt so far?

In the preceding sections, we presented evidence for the existence of muscle gearing and its underlying mechanisms predominantly from animal studies and computational models. A common question in the biological sciences is whether the physiological and mechanical behaviours observed in animal studies can be extrapolated to living animals (including humans) operating with muscles functioning *in vivo* under normal physiological conditions. This section synthesises current *in vivo* research on muscle gearing and highlights potential commonalities and discrepancies between animal and human studies.

A summary of studies investigating muscle gearing is presented in Table 2 and includes *in situ*, *in vitro, ex vivo*, and *in vivo* investigations across both non-mammalian and mammalian pennate muscles. Reported gear values, operation ranges, and responses to mechanical demands vary across experimental models. In isolated muscle preparations from animals, gear ratios typically range between ∼1.0 and 2.0 during singular concentric contractions and from 1.5 and 4.5 during singular eccentric contractions, i.e., when concentric and eccentric contractions are not performed consecutively like in a stretch-shorten cycle. In contrast, *in vivo* human studies suggest gear ratios between 1.0 to 1.5 for concentric contractions and approximately 1.0 to 3.1 for eccentric contractions.

**Table 2.**
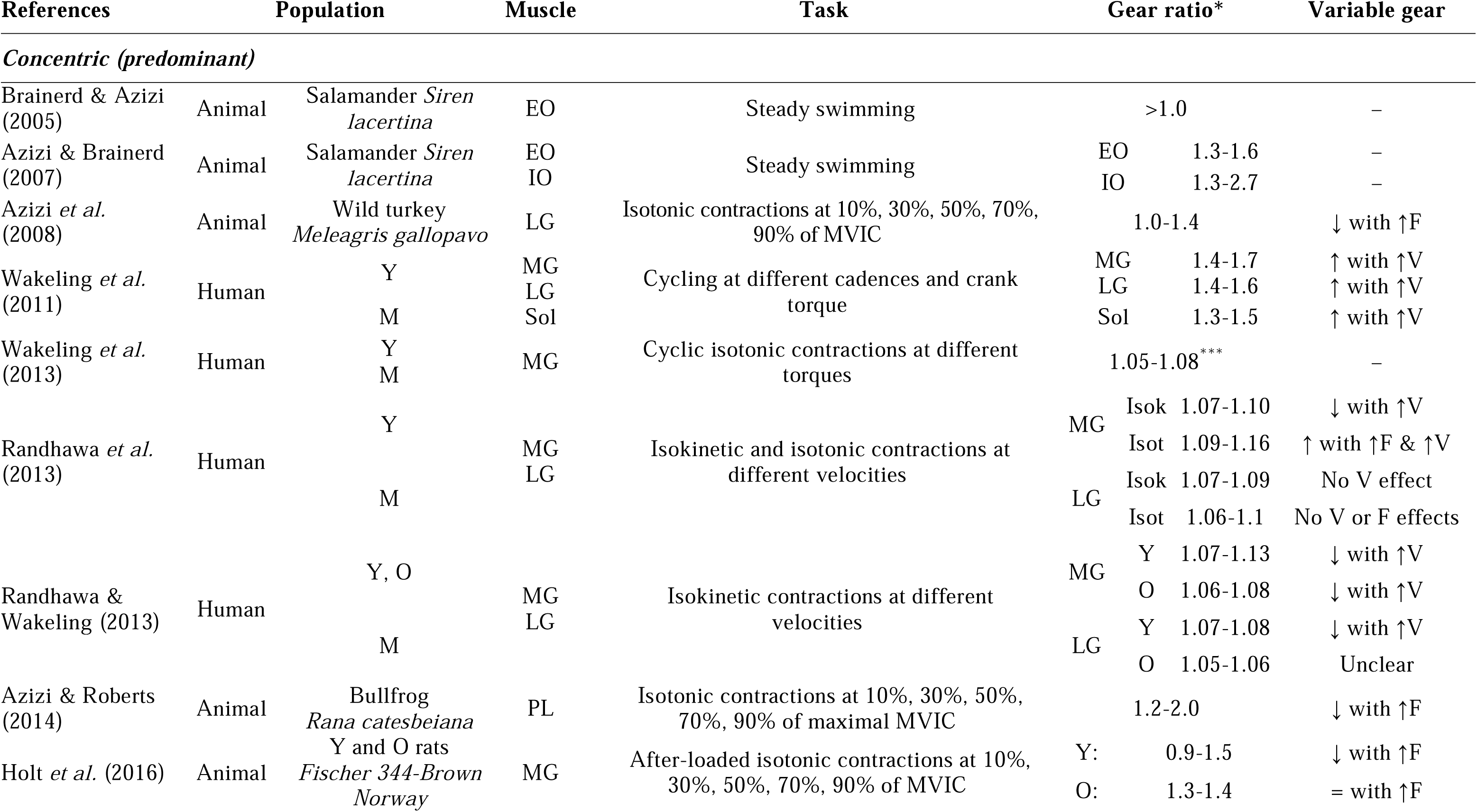
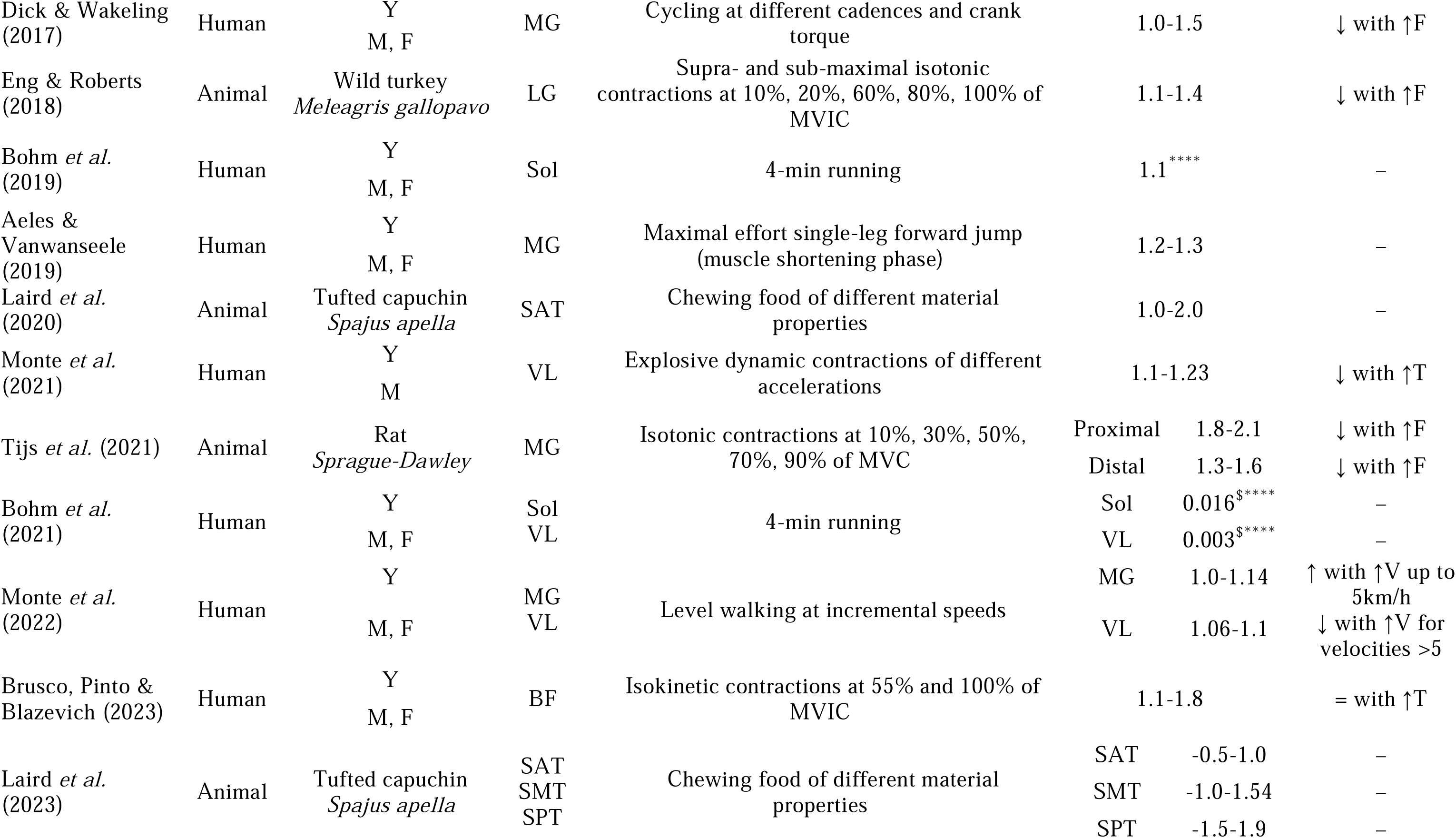
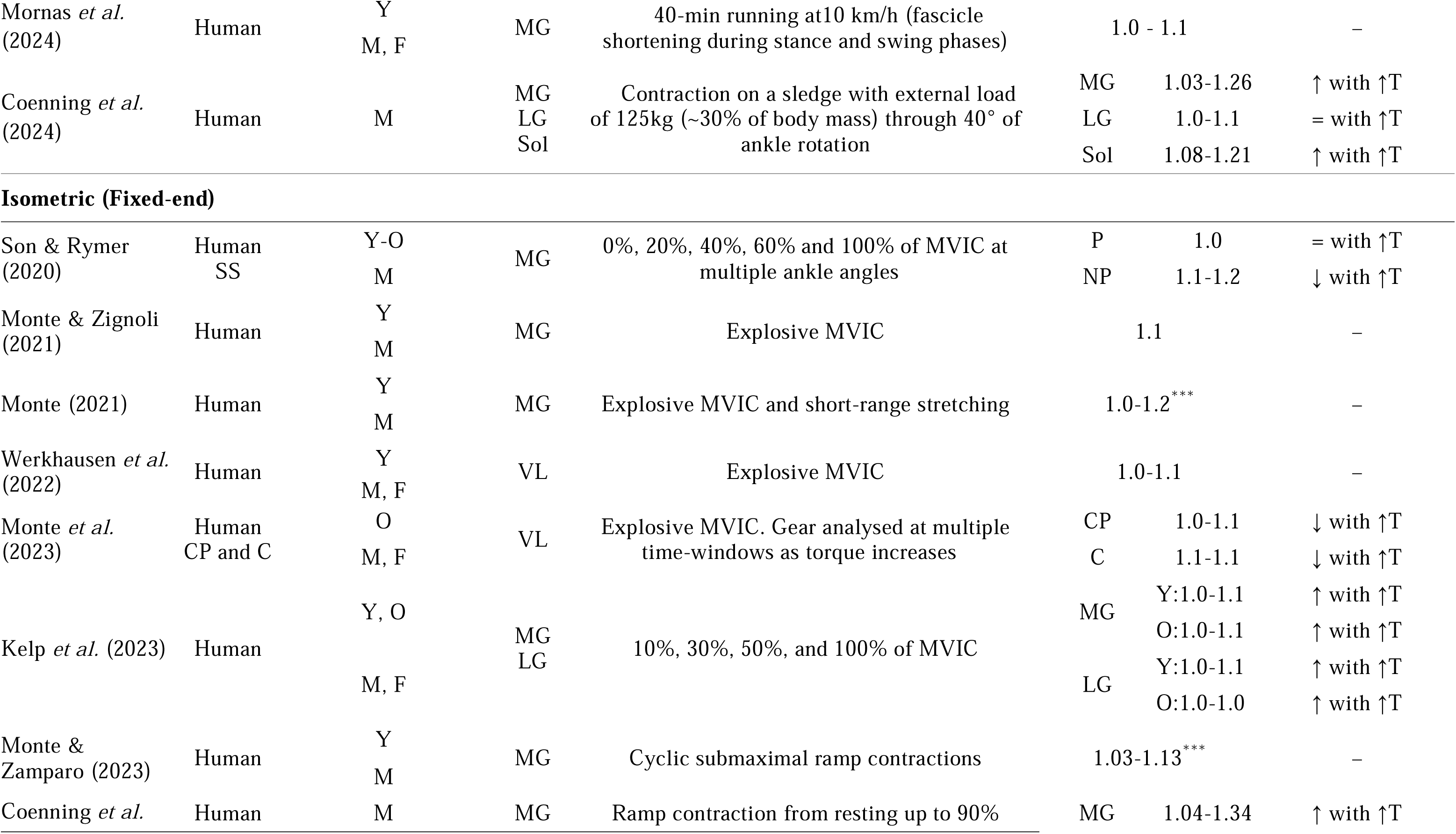
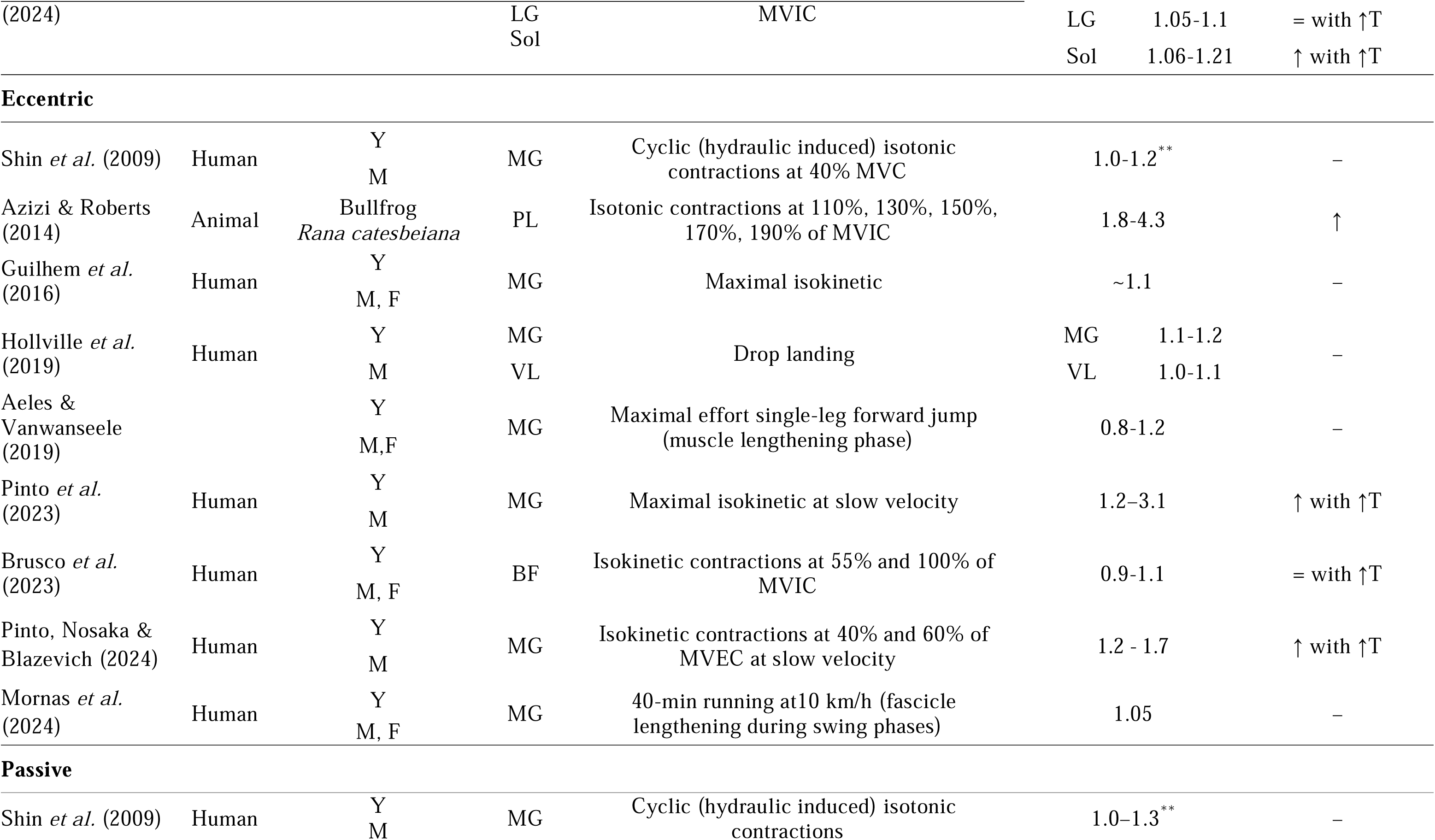
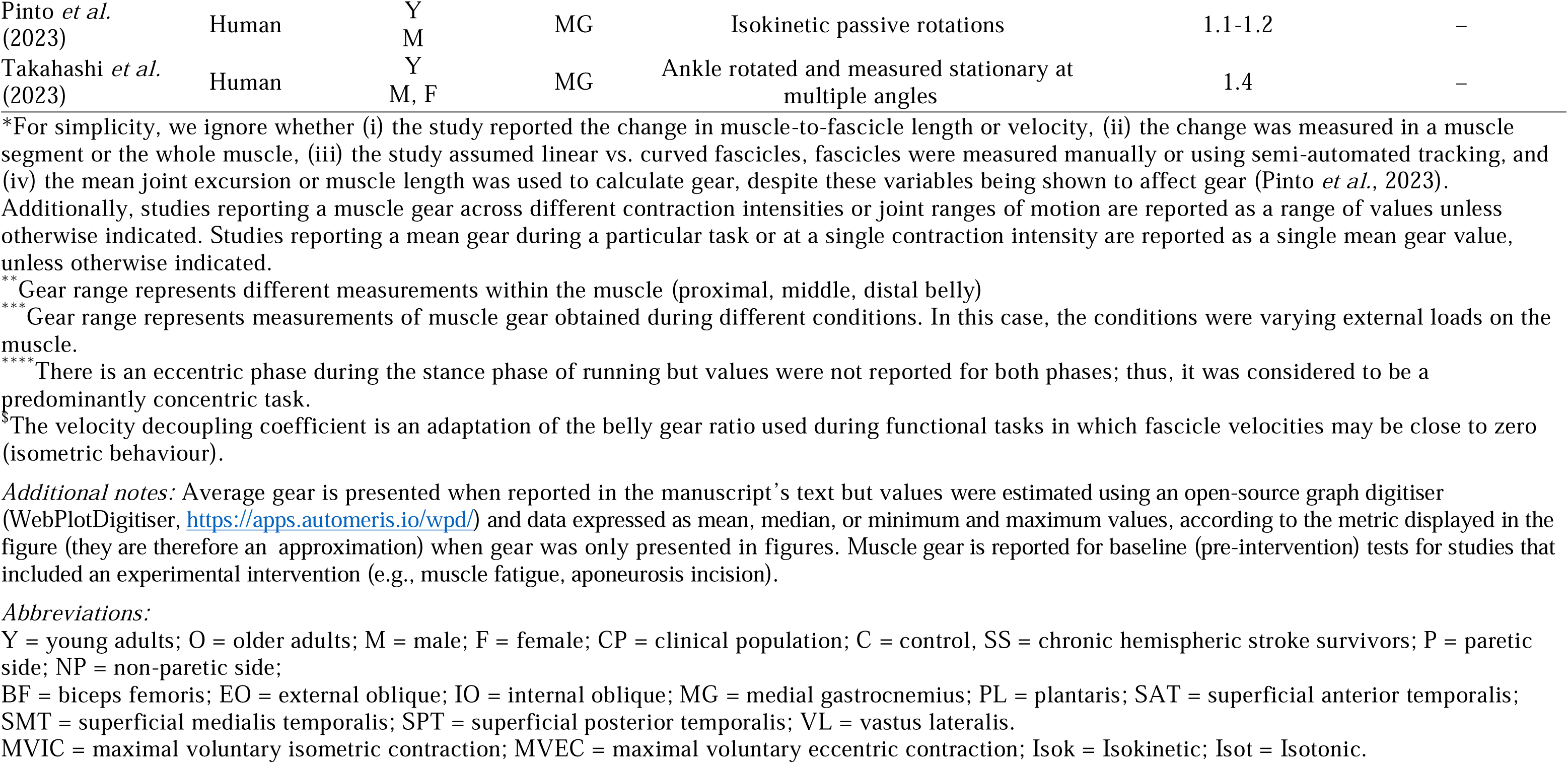
Summary of studies reporting muscle gearing during concentric, isometric, and eccentric contractions as well as passive lengthening.

Both animal and human studies have reported torque- or force-dependent shifts in gear ratios, although only studies in humans have reported gear to vary with velocity. Furthermore, differences in variable gearing exist, with most *in vivo* human studies reporting gear ratio to increase with force or torque (or to increase with velocity) (Son & Rymer, 2020; Kelp *et al*., 2023; Randhawa *et al*., 2013; Coenning *et al*., 2024) whereas animal studies report gear to decrease with force increase. To our knowledge, no studies on animals muscles have demonstrated an increase in gear ratios with force, and only two studies in humans have reported a gearLforce (or torque) relationship similar to that of animal experiments (Dick & Wakeling, 2017; Monte *et al*., 2021).

The reasons for these discrepancies remain unclear and may arise from variations in experimental approaches – *in situ*, *in vitro*, or *ex vivo* versus *in vivo* human studies – or they may reflect fundamental differences in muscle function between human and non-mammalian animals.

### (1) Differences in testing paradigms

Experimental approaches for testing muscle gearing differ substantially between animal and human studies, resulting in considerable variations in the mechanical conditions tested. Methodological discrepancies may influence observed outcomes and interpretations of how force and/or velocity modulate muscle gearing.

In animal studies, muscle gearing is examined under highly controlled laboratory conditions using servomotor-driven experiments that enable force and length measurements. The muscle-tendon unit is surgically exposed and freed from surrounding muscles, with one tendon and its attached bone fixed to a rigid apparatus while the other tendon is calcified and clamped to a servomotor lever arm (Azizi & Roberts, 2014; Azizi *et al*., 2008). Fascicle lengths and velocities are estimated using sonomicrometry (surgically implanted piezoelectric crystals) while whole□muscle length and velocities are measured via the servomotor. Electrical stimulation via implanted cuff electrodes to branching nerves induces tetanic contractions against predefined forces or muscle lengths.

A common protocol used for gearing assessment begins with an isometric ramp contraction, during which force is gradually increased to a predefined percentage of the maximal isometric force. This is followed by an isotonic phase, where muscle force is held constant as the muscle (and its fascicles) shortens or lengthens at approximately constant velocities. Gear ratio is measured during this constant□force phase to minimise the influence of series elastic element compliance (Azizi *et al*., 2008), as tendon elasticity affects muscle length changes (discussed below). A key advantage in these experiments is the ability to control initial fibre length and pennation angle, which are kept the same at the onset of contractions across force levels, including during the initial plateau of the isometric phase (see Figure 3 in Azizi *et al*., 2008). Furthermore, gear ratios are determined during the constant-force phase occurring at or near the plateau of the force−length relationship at the muscle’s optimal fascicle (L_0_) during all force conditions (Azizi *et al*., 2008; Holt *et al*., 2016). These controlled testing conditions ensure that gearing is not confounded by differences in fascicle strain, force−length properties, or initial pennation angle (Brainerd & Azizi, 2005), while also minimising variations in fascicle radial expansion due to passive elastic properties. This effectively isolates muscle shape changes as the primary driver of gearing and enables comparisons between muscles to determine whether differences in gear arises from variations in muscle shape changes or in initial pennation angles.

In contrast, human *in vivo* studies rely on non-invasive imaging techniques, such as ultrasound, to estimate fascicle length and muscle-tendon junction displacement (Roberts & Dick, 2023). When ultrasonography is combined with motion analysis systems, accurate estimations of whole-muscle length changes are possible. While conceptually similar to the animal preparation experiments that use sonomicrometry and servomotors, human experiments lack the precise control over muscle forces, velocities, and activation timing. Such limitations make direct comparisons with *in vitro* and *in situ* animal studies challenging. A variety of experimental paradigms have been employed in human studies (see Table 2), including isometric, isotonic, and isokinetic contractions, as well as more complex tasks involving time-varying force and velocity (Monte *et al*., 2022). Muscle gearing has been investigated during isometric contractions under both time□varying (explosive) (Monte *et al*., 2023; Monte & Zignoli, 2021) and approximately constant force or torque conditions (Kelp *et al*., 2023; Monte, 2021). Dynamic contractions have been tested, such as isotonic (Randhawa & Wakeling, 2013; Randhawa *et al*., 2013), explosive (Monte *et al*., 2021), and isokinetic (Randhawa & Wakeling, 2013; Randhawa *et al*., 2013; Pinto *et al*., 2024; Pinto *et al*., 2023) contractions. In these studies, external loads, cycle frequencies, and set velocities have been systematically manipulated to assess their effects on muscle gearing (Randhawa & Wakeling, 2013; Randhawa *et al*., 2013). Some studies have used cycling-based paradigms, where bicycle crank torque and velocity can be manipulated via bicycle gear ratios and cadences to partly decoupling of force- and velocity□effects on muscle gearing (Wakeling *et al*., 2011; Dick & Wakeling, 2017). However, unlike in constant-force paradigms in animal models, many of these human studies determine gear ratios during force-changing conditions and estimating muscle segment velocities (Wakeling *et al*., 2011; Monte *et al*., 2021), which can alter interpretation of findings (Pinto *et al*., 2023). Other methodological differences also exist. For example, gear ratios have been calculated during transient (Dick & Wakeling, 2017) or time-averaged periods (Pinto *et al*., 2024; Pinto *et al*., 2023; Monte *et al*., 2023; Monte *et al*., 2022). Gear ratios have been assessed at specific time points during the contraction cycle (Monte *et al*., 2022), at different parts of the contractile cycle during peak muscle and fascicle velocities (Randhawa & Wakeling, 2013; Randhawa *et al*., 2013; Dick & Wakeling, 2017), or at the muscle’s optimal fascicle length (Monte *et al*., 2021). While findings mostly show variability in gearing, results suggest that gear ratios depend on force or velocity, with force-dependent gearing often, though not always, similar to that observed in animal studies where gear decreases with force (see Table 2).

The significant methodological differences and testing paradigms between animal and human complicate direct comparisons of how force and velocity affect muscle gearing. The precise control over force, velocity, and starting fascicle angles and lengths in isolated animal muscle experiments is difficult to replicate *in vivo*. However, studies examining rat medial gastrocnemius shape changes *in vivo* under time-varying force conditions, such as during gait and slope conditions like decline walking or incline galloping, have shown force-dependent shape changes similar to those observed in animal models (Konow, Collias & Biewener, 2020). This raises the question of whether observed differences in muscle gearing primarily stem from experimental design and muscle preparation or reflect inherent functional differences between humans and other animals

The muscle model presented herein addresses some of these challenges by incorporating testing paradigm similar to those used in animal studies while also simulating activation levels comparable to human experiments. The model demonstrates that gearing is independently affected by force, activation, and velocity, with high gear ratios occurring at high velocities independent of muscle force. Of note, this model does not include an in-series tendon, meaning that tendon compliance effects could not be a contributing factor to observed gearing behaviours in the simulation. Thus, differences in testing paradigms should be carefully considered when interpreting muscle gearing results and extrapolating findings from animal models to human movement. Nonetheless, potential intrinsic species differences should not be overlooked (see below).

Given the substantial variability in human protocols, and the tendency for force, activation and shortening velocity to covary, future studies should aim to decouple these effects. Whilst it is difficult to measure muscle force in humans (though estimates are possible using musculoskeletal modelling and inverse dynamics), one approach would be to systematically manipulate load (e.g., isotonic contractions), prescribed velocity (e.g., isokinetic contractions), and activation levels (quantified volitionally or controlled via electrical stimulation), while quantifying time-varying whole-muscle and fascicle velocities from which instantaneous, time-resolved gear ratios are derived. Where the aim is mechanistic inference, gearing should be evaluated at comparable operating lengths (near L_0_) and under clearly defined force–velocity–activation states; however, in task-based contexts (e.g., locomotion), the priority may instead be to characterise how gearing emerges under naturally time-varying forces, velocities, and activations. More broadly, protocols that incorporate constant-force/constant-torque phase or factorial combinations of load and velocity, and activation would improve comparability across studies and species.

### (2) Factors within the muscle-tendon unit (MTU)

In animal experiments, muscle gearing is measured during conditions where force is constant and the influence of the tendon is minimised (Azizi & Roberts, 2014; Holt *et al*., 2016; Eng & Roberts, 2018; Azizi *et al*., 2008). To achieve this, the in-series tendon is either calcified (Azizi *et al*., 2008) or otherwise made short and stiff by clamping it close to the muscle’s end (Tijs *et al*., 2021; Holt *et al*., 2016). These methodological constraints, combined with the constant-force condition, limit the contribution of the in-series tendon to force□dependent muscle length changes. As a result, muscle shape changes and gearing occur with minimal interference from tendon elasticity. However, it is nearly impossible to remove the influence of tendon compliance in human muscles with long tendons, such as the gastrocnemii. Tendon elasticity significantly affects muscle force−length and force−velocity relationships as well as overall mechanics of the muscle-tendon unit. During contraction, muscles fibres shorten and increase in pennation, while the in-series tendon stretches to accommodate these changes. This tendon compliance influences initial pennation angles, which can in turn influence gearing (Brainerd & Azizi, 2005). Unlike in animal experiments, human studies cannot precisely control for these effects. Consequently, tendon elasticity may cause muscle gearing to increase rather than decrease with force, leading to different outcomes than those observed in isolated muscle preparations.

The mechanical properties of a muscle’s ECM may impact shape changes during contraction and therefore gearing (Eng *et al*., 2018; Roberts *et al*., 2019). Whilst our model shows variable gearing despite absence of an aponeurosis or specific connective tissues around the muscle, ECM components may still contribute to muscle shape changes and gearing. Muscle fibres and fascicles are encapsuled by endomysial and perimysial connective sheets, and research has shown that their quantity and morphology vary among vertebrates (Sleboda *et al*., 2020) and even among muscles within the same species (Purslow, 2020). Given that ECM properties appear to influence shape changes at the fibre and fascicle levels, one might expect gearing differences to exist between functionally similar muscles across species, as well as between different muscles within the same species. However, there is limited evidence to determine whether, or the extent to which, between-species ECM structural and mechanical property variations can explain differences in gearing between animals and human muscles. This is an area that would benefit from computational modelling studies where the effect of tissue manipulations can be addressed directly.

Spatial variations in motor unit activations also occur during tasks with different force or velocity characteristics (Wakeling, 2009) and influence muscle metabolic cost (Lai, Biewener & Wakeling, 2018) and muscle force (Wakeling *et al*., 2012) possibly through alterations in fascicle behaviour and gearing. Rahemi *et al*. (2014) developed a 3-D finite element model based on human lateral gastrocnemius to explore the effect of submaximal isometric contractions with uniform and non-uniform (regionalised) activation on fibre and muscle behaviours. All models had 10% of the fibres activated. Regionalised activation also caused greater regionalisation of thickness changes, resulting in larger fascicle strains and greater changes in pennation angle. These findings suggest that regionalised activation can influence shape changes and thus variable gearing (Rahemi *et al*., 2014). Differences in spatial fibre activation should affect muscle shape, particularly during submaximal activation, as regions of inactive fibres do not contribute to force production but still add mass, increasing non-active tissue inertia (Holt, Wakeling & Biewener, 2014; Ross *et al*., 2020) and potentially limiting radial bulging of active fibres and fascicles. These may in turn reduce contraction efficiency (Ross & Wakeling, 2021; Ross *et al*., 2020) through its impact on gearing.

Spatial activation variation is an inherent physiological phenomenon also reported in animal muscles *in vivo* (Higham & Biewener, 2011; Higham, Biewener & Wakeling, 2008). However, studies examining variable gearing in isolated muscle preparations are conducted under supra-maximal activation conditions in which the whole muscle is assumed to be uniformly activated (Eng & Roberts, 2018). An exception to this was shown in the isolated rat medial gastrocnemius (Tijs *et al*., 2021) in which lower gear ratios were observed during contractions triggered by submaximal rather than maximal stimulation intensities, although it did not reduce fibre shortening velocity or alter the gear−force relationship (Tijs *et al*., 2021). It remains to be determined whether these findings are applicable to muscles functioning *in vivo* due to differences between asynchronous voluntary (as in *in vivo*) vs. synchronous electrically-evoked motor unit stimulation patterns.

As with some animal muscles, regional muscle architectural variation is also observed in human muscles (Takahashi *et al*., 2022; Shin *et al*., 2009) and has the potential to affect gearing (Tijs *et al*., 2021; Pinto *et al*., 2023; Azizi & Deslauriers, 2014; Takahashi *et al*., 2023; Shin *et al*., 2009). Regional pennation variation can result in asymmetric transverse muscle belly strains, regional fascicle reorientations, fascicle velocities, and muscle segment length changes (Blazevich, Gill & Zhou, 2006). Most gearing studies are conducted in small animals possessing relatively small muscles and relatively homogenous architecture (Azizi & Roberts, 2014; Eng & Roberts, 2018), although this is not always the case (Tijs *et al*., 2021; Holt *et al*., 2016). Recently, Tijs *et al*. (2021) demonstrated region-specific gear ratios in the rat medial gastrocnemius, with lower gear in muscle compartments exhibiting lower pennation regardless of activation status. However, gear changes with contractile force were similar to those reported in previous small animal data sets.

Finally, intramuscular adipose tissue (fat) may be a potential factor influencing variable gearing (Rahemi *et al*., 2015; Kelp *et al*., 2023). Fat accumulates within skeletal muscle tissue (e.g. with ageing and disuse) and can be stored around myofibrils (intramyocellular fat or lipids droplets) and surrounding muscle fibres and fascicles (extramyocellular fat) (Altajar & Baffy, 2020). Muscle must use internal work to deform the fatty inclusions, and this affects the way in which the contractile tissue deforms (Wakeling et al. 2020). Fat may increase the stiffness of the muscle and this increase would resist fibre shortening and transverse bulging of fibres and the muscle belly, ultimately reducing fascicle reorientation and gear (Rahemi *et al*., 2015; Kelp *et al*., 2023). Determining fat quantities in animal studies would help pinpointing whether fat could be a contributing factor to the gearing differences previously observed between humans and other, non-mammalian animals.

### (3) Factors residing outside the muscle

In many vertebrates, joint movement is produced by the activation and subsequent length change of multiple muscles tightly packed in muscle groups (compartments). Unlike studies using animal preparations that typically isolate muscles from others within their group to measure gearing, human muscles normally operate within their synergistic group *in vivo* (Reinhardt *et al*., 2016; Bernabei *et al*., 2017). Thus, gearing can be influenced by external factors constraining radial expansion during muscle shortening, including compressive loads exerted onto a muscle by bones, inter-muscular fat, or neighbouring muscles within the group.

Several factors can influence lateral muscle pressures and deformations in synergistic muscle groups *in vivo*. First, synergistic muscles may lack common neural drive (Hug *et al*., 2021), resulting in varying activation levels among muscles of a group. Second, synergistic muscles may possess different shapes and architectures (Aeles *et al*., 2022; Blazevich *et al*., 2006) and deform asymmetrically as they shorten and expand radially. Asymmetric radial expansion in a muscle within the group may promote lateral pressures differences along adjacent muscles (Bernabei *et al*., 2017). Third, individual muscles within the group may attach proximally and distally through distinct tendons or (intermuscular) aponeuroses that differ in their mechanical properties (Bernabei *et al*., 2017; Ando *et al*., 2018), so muscle shortening (or lengthening) during contraction between muscles may differ even if they exert similar forces. Fourth, the presence of non-contractile materials such as inter-muscular fat and connective tissues may influence muscle deformations. And finally, differences in architectural features imply distinct muscle force contributions to the total synergistic force or joint torque (Wakeling *et al*., 2011; Higham & Biewener, 2011). While these factors may be inter-related, their independent or combined effects may impact muscle length change and radial pressures even when they are subjected to the same external loads, and their effects on fascicle reorientation and gearing might not be easily or accurately estimated using current muscle models.

The first evidence demonstrating an effect of external force constraining radial bulging of human muscle and fascicle behaviours *in vivo* came from studies in which external compression was applied using elastic bandage. Wakeling *et al*. (2013) found reductions in belly gear, fascicle angle, and muscle thickness changes during contraction without significant effect on concentric joint torque production. Under these conditions, belly gear reductions imply an increased fascicle shortening velocity that would decrease fascicle force-generating potential. Other studies have since imposed transverse pressures using external loads or compressive garments to compress the muscle and observed a significant impact on muscle geometry (Ryan *et al*., 2019; Stutzig *et al*., 2019; Monte, 2021), force production (Ryan *et al*., 2019; Stutzig *et al*., 2019; Siebert *et al*., 2018), and energetic cost (Monte & Zamparo, 2023), with both empirical data from animal studies and finite element modelling indicating that compression-related force reduction is exacerbated when a muscle is at shorter length (Sleboda & Roberts, 2020) or has lower pennation (Ryan *et al*., 2020a). These effects appear to be direction dependent, with loads placed in the thickness direction increasing longitudinal muscle forces at muscle lengths shorter than resting and decreasing for shorter lengths, whereas loads compressing in the width direction promote reductions in longitudinal forces without length-dependency effects (Ryan *et al*., 2020a). When muscle is compressed from all directions and at shorter muscle lengths, no changes in force or fascicle behaviour are detected (Ryan *et al*., 2020b)

Evidence indicating that architectural differences between synergistic muscles can contribute to relative lateral pressure variations has been partially presented. Randhawa *et al*. (2013) showed that MG reduced in thickness during both isovelocity and isotonic concentric plantar flexion whereas the LG increased thickness. These observations were subsequently replicated in other studies (Randhawa & Wakeling, 2018; Wakeling *et al*., 2011; Kelp *et al*., 2021; Kelp *et al*., 2023) and demonstrated that between-muscle synergism occurs such that as one muscle reduces in thickness the other may increase in response to the pressure reduction in that direction (to ‘fill the space’). This interaction, in which adjacent muscles exert pressure and perform work on each other, likely influences muscle shape changes. Such effects are not accounted for in animal experiments and may contribute to differences in muscle gearing between human and animal preparations.

## VII. Other functional benefits of fibre arrangement and muscle shape changes during contraction

Muscle shape changes during contraction may confer additional benefits beyond their effects on fascicle reorientation and muscle gearing. Because muscles operate within tightly packed anatomical compartments, transverse expansion during contraction can lead to compression of adjacent structures and reduce mechanical efficiency (Wakeling *et al*., 2020). Muscle architecture and shape changes may be finely tuned not only for mechanical efficiency but also for spatial accommodation. Features such as fibre orientation, muscle belly shape and curvature, and the direction of force transmission can influence how a muscle deforms during contraction, enabling it to avoid impinging surroundings. Dynamic changes in muscle shape offer an elegant anatomical solution for managing spatial constraints while preserving the functional and anatomical integrity of both the muscle and surrounding tissues.

One example of this comes from the adductor mandibulae of *Creatochanes*, a teleost fish with big eyes (Alexander, 1964, 1983). In this species, the main jaw-closing muscle exhibits a crescent-like, concave shape that partially wraps around the posterior and ventral surfaces of the eyeball (Alexander, 1964). If this muscle had a parallel-fibred architecture, contraction would cause it to bulge radially and compress the adjacent orbital tissue. If the muscle had a pennate configuration, it could be designed in a way that contraction would cause muscle expansion predominantly in the opposite direction of the eye, avoiding compression of ocular structures. In reality, the adductor mandibulae muscles exhibit a pennate architecture with fibres inserting obliquely onto a central subocular tendon lying along the inner concave edge of the muscle, adjacent to the eye. Thus, the contracting fibres pull the tendon away from the eyeball without requiring the muscle belly to expand toward the orbit (Alexander, 1983; Datovo & Vari, 2013). In effect, this muscle architecture offers both a mechanical and spatial solution to optimise force production while protecting sensitive surrounding structures from compression.

A comparable example in humans involves the MG and LG, which act synergistically during plantarflexion. These muscles differ in shape, fibre orientation, and deformation patterns (Randhawa & Wakeling, 2018; Wakeling *et al*., 2011; Kelp *et al*., 2021; Kelp *et al*., 2023). While the MG tends to reduce in thickness during contraction, the LG increases in thickness. This reciprocal deformation pattern may represent a spatially effective and energetically favourable strategy in which the muscles coordinate shape changes to avoid competing for space and generating unnecessary internal work to deform each other, thus maximising net external output. Overall, these examples support the broader hypothesis that muscle architecture, and its associated muscle shape changes during contraction, has evolved to optimise both mechanical performance and anatomical compatibility within tightly constrained anatomical environments. Understanding these interactions could inform the design of bioinspired actuators or therapeutic strategies that consider both force output and spatial constraints. Nonetheless, further research is warranted to fully understand the trade-offs involved.

## VIII. Clinical and practical implications

There are several benefits to improving our understanding of muscle gearing within the clinical context, including potentially gaining insights into muscle functional loss with ageing, neurological disorders, and physical training and detraining.

Ageing is typically accompanied by (i) reductions in muscle strength and power (Freitas *et al*., 2024), (ii) a reduction in resting pennation angle (Morse *et al*., 2005; Narici *et al*., 2003; Tomlinson *et al*., 2014; Randhawa & Wakeling, 2013), (iii) increased longitudinal and transverse ECM stiffness resulting from alterations in ECM geometry and increased cross-linking (Wood *et al*., 2014; Kragstrup, Kjaer & Mackey, 2011), (iv) increased intra-and inter-muscular fat content (Addison *et al*., 2014; Kelp *et al*., 2023), (v) reduced intramuscular fluid with reductions in intracellular and maintenance of extracellular water (Yamada *et al*., 2010; Iwasaka *et al*., 2023), and (vi) irregular myofiber shape (Soendenbroe *et al*., 2023), and (vii) fibre type loss, motor unit remodelling, and regional heterogeneity in fibre type distribution (Jones *et al*., 2022). Alterations in one or more of these structural factors have the potential to influence gearing. Growing evidence suggests that muscle gearing is influenced by biological age (Holt *et al*., 2016; Randhawa & Wakeling, 2013; Kelp *et al*., 2023), or at least the increase in inactivity that comes with ageing. In both rats (Holt *et al*., 2016) and humans (Randhawa & Wakeling, 2013), older muscles have been found not to change gear as much in response to load (Holt *et al*., 2016) or velocity (Randhawa & Wakeling, 2013) changes. The inability to change gear in response to a task’s mechanical demands (e.g., force or velocity) may at least partly explain the age-related decline in both muscle strength and power. As illustrated in Figure 11, the age-related difference in *fibre* shortening peak power is smaller than the difference in *whole muscle* peak power. Furthermore, and perhaps of greater functional importance, there is substantial narrowing of the *whole muscle* power−velocity relation due to the inability of muscles to vary gear in response to changes in load (see Figure 3 and 11B). These results were observed in aged rats and are similar to those demonstrated in single fibre preparations from human muscles (Grosicki, Zepeda & Sundberg, 2022) as well as whole muscle peak power and width of the power−velocity relation *in vivo* (Alcazar *et al*., 2023). Thus, alterations in variable gearing capacity offers a promising mechanistic explanation for some of the age-related decline in the power and strength in older adults, although further studies in humans are necessary to test this hypothesis.

**Figure 11.**
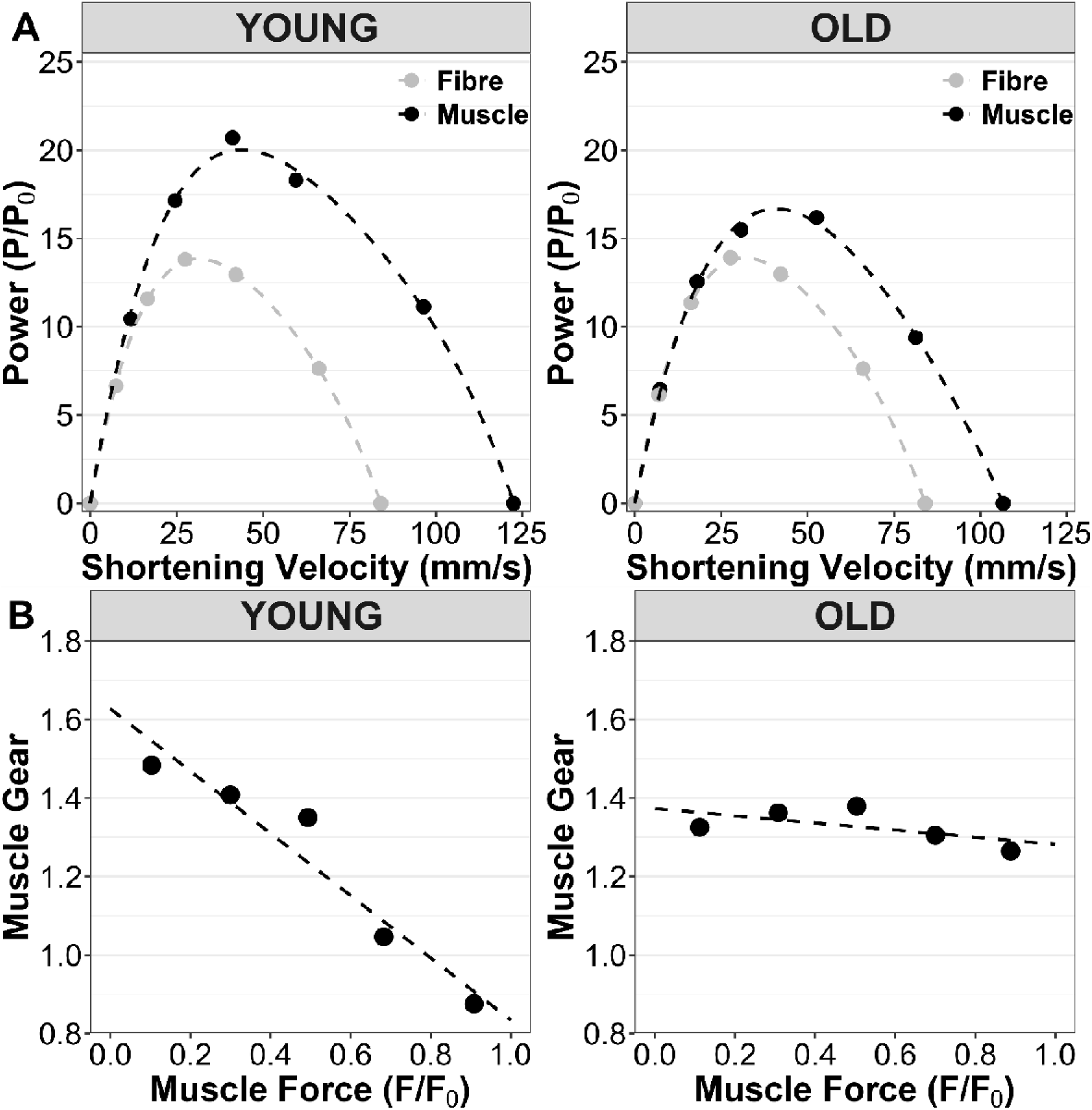
(A) Differences in fibre and whole muscle power-velocity relation in young and old rats. (B) Muscle gear in young (left) and old (right) rats. Gearing varies with force in young but not old rats. Old rats work at similar gears in the high force and low-force contractions, and this is hypothesised to be affected by increases in stiffness of both aponeurosis and the extracellular matrix surrounding fibres and fascicles. Figure adapted from Holt et al. (2016). Power was calculated from the available shortening velocity and whole muscle force (normalised to peak isometric force). Values were estimated using an open available graph digitiser (WebPlotDigitiser, https://apps.automeris.io/wpd/).

Second, several musculoskeletal disorders as well as recovery from muscle injury are associated with substantial alterations in the quantity and structure of the ECM and intramuscular fat. For example, increased collagen content (fibrosis), altered collagen architecture, and increased collagen cross-linking have been observed in Duchenne muscular dystrophy (Wohlgemuth *et al*., 2023; Brashear *et al*., 2021) and cerebral palsy (Smith *et al*., 2011b; Lieber & Fridén, 2019) as well as following muscle strain injuries (Reurink *et al*., 2015) and both ligament tears and reconstruction (Noehren *et al*., 2016; Peck *et al*., 2019). In contrast, classic-like Ehlers□Danlos Syndrome, a genetic connective tissue disorder caused by tenascin□X deficiency in the ECM, leads to reduced collagen fibril density and length in the endomysium and more compliant connective tissue within and between muscles (Huijing *et al*., 2010; Voermans *et al*., 2007). Disease-related alterations in ECM architecture and mechanical properties have the potential to alter intermuscular myofascial force transmission between neighbouring muscles (Huijing, 2003; Huijing *et al*., 2010), thereby changing compartment-level external constraints on muscle shape change and fascicle reorientation. Accumulation of intra- and inter-muscular fat content are also a common response to skeletal muscle injury and disease (D’Souza *et al*., 2020; Akinci D’Antonoli *et al*., 2021; Leroy-Willig *et al*., 1997), with both persistent fatty infiltration in the interfascicular space and within myofibrils even after a short□term 3□month resistance training (Bayer *et al*., 2021). The internal work needed to deform these non-contractile tissues may detract from the external work that the muscle can do (Rahemi *et al*., 2015; Wakeling *et al*., 2020; Konno *et al*., 2021b) and alter the way the muscle fascicles can change in length and pennation. It is therefore possible that the persistent muscle strength, speed reductions, or even altered metabolic cost and muscle efficiency after injury and disease may be influenced by altered muscle gearing.

Third, changes in muscle gearing could play a role in the changes in voluntary and involuntary (electrically□induced) contractile capacity in response to exercise-based warm-up, which has been shown to enhance performance (Blazevich & Babault, 2019). This is because brief bouts of muscle contraction can trigger substantial fluid shifts into the muscle (Sjøgaard, Adams & Saltin, 1985; Ploutz-Snyder, Convertino & Dudley, 1995), which should then enhance contractile capacity by (i) affecting the interactions between collagenous ECM tissues and the fluid to increase muscle stiffness (Sleboda & Roberts, 2017; Sleboda *et al*., 2019), (ii) increase intramuscular pressure, (iii) increase resting pennation and subsequent reorientation, and (iv) possibly reduce the distance between each contractile unit (fibres and fascicles), which all impact gearing. It has been hypothesised that post□warm□up fluid shifts should increase muscle gear and benefit fascicle force−velocity and force−length properties during contraction (Blazevich & Babault, 2019). Indeed, Bourgeois, Duchateau & Baudry (2022) recently demonstrated small yet significant increases in peak twitch and doublet forces accompanied by increases in medial gastrocnemius fascicle reorientation and shortening. These changes occurred in the absence of detectable alterations in resting architecture and are unlikely to be attributed to changes in series elastic compliance (Gago *et al*., 2014) after performing the conditioning 6□s maximal voluntary contraction. Such finding is consistent with an acute contraction-induced alteration in gearing.

Lastly, many physical exercise interventions, whether combined with nutritional strategies or not, often immediately reduce (fatigue) or enhance (potentiation) muscle force. Some studies also report post□exercise changes in muscle fluid (Sjøgaard *et al*., 1985; Ploutz-Snyder *et al*., 1995; Beis *et al*., 2011; Antonio *et al*., 2021; Ribeiro *et al*., 2020; Francaux & Poortmans, 1999), ECM remodelling (Csapo, Gumpenberger & Wessner, 2020; Guzzoni *et al*., 2018), intra□ and inter□muscular fat (Langleite *et al*., 2016; Ramírez-Vélez *et al*., 2021), and resting pennation angle (Kawakami *et al*., 1995; Timmins *et al*., 2016), which may all influence muscle gearing. Nonetheless, the extent to which physical exercise interventions acutely or chronically affect muscle gearing remains uncertain, and it is unclear whether the observed changes in muscle force and function following these interventions can be attributed to alterations in muscle gearing.

## IX. Conclusions

1. Skeletal muscles exhibit a broad range of contractile behaviours that are strongly influenced by their internal architecture. Historically, our understanding of the structure−function relation has been largely derived from muscle structure measurements obtained at rest, through anatomical dissections or imaging techniques, and its function determined during isometric or concentric contractions. However, muscles typically operate under dynamic conditions involving both active and passive shortening and lengthening, during which substantial structural and shape changes occur. These structural deformations critically influence the most fundamental properties of muscle contraction (i.e., force−velocity and force−length relationships) and cannot be accurately predicted from examination of resting (non-contracting) architectural features alone.
2. Structural changes during muscle contraction have been documented for over a century (since at least the 1920’s), yet only recently has the functional significance of changes in fascicle angle and its influence on muscle shape changes during contraction been recognised. Fascicle reorientation serves as a gearing mechanism, partially decoupling fascicle velocity from whole-muscle velocity and thereby providing a mechanical (velocity) advantage. Gearing enables muscle fascicles to operate closer to their optimal force−length and force−velocity relations while facilitating greater whole-muscle level length change or contraction velocity. Importantly, in most cases the extent of this decoupling is not fixed; it varies dynamically with the mechanical demands of the task, allowing muscles to modulate gear ratios in real-time, effectively broadening their functional range and mechanical efficiency.
3. Recent studies have investigated the mechanisms underpinning variable gearing, most often using bioinspired muscle models or isolated muscle preparations subject to supramaximal, short-duration electrical stimulations. These investigations have examined factors including the roles of pennation angle, directional components of contractile elements (fibres) forces, three□dimensional muscle expansion, intrinsic and extrinsic tissue properties, intramuscular pressures, external pressures, and mechanical work on the muscle surface. Variable gearing is a multifactorial phenomenon but yet appears to be an emergent property of muscle function, and emerges from the complex, non□linear interactions among all these factors.
4. The computational modelling introduced herein integrates these mechanisms and demonstrates that the general features of muscle gearing emerge from the physical properties of the contractile elements and the muscle tissue properties alone, without requiring extrinsic structures such as aponeurosis or other connective tissue structures. The simulations reproduced the shape changes and gearing effects reported in maximally activated animal muscles and extended these predictions to more nuanced behaviours observed under submaximal activation levels typical of human movement. The modelling framework showed that force, velocity, and activation level independently modulate muscle gearing, with the most pronounced effects occurring at higher contraction velocities, where reducing fibre shortening velocity can enhance force output.
5. Real-time changes in muscle gear in response to contractile conditions have been documented in both human and animal pennate muscles. While many animal studies suggest a decrease in gear with increasing force, human *in vivo* studies have shown more variable responses of gear changes, including increases with force or velocity, decreases with force or torque, or no change at all. These discrepancies may reflect both methodological and biological differences, such as whether muscle force is held constant during measurement, the mechanical influence of compliant tendons, and the complex deformation patterns associated with synergistic muscle behaviour *in vivo*; such conditions differ markedly from those of controlled animal experiments.
6. Muscle architecture and shape changes during contraction may offer function al advantages beyond influencing variable gearing. Because muscles operate in confined anatomical spaces, their deformation patterns may be tuned not only for force or velocity production but to prevent interference with adjacent tissues. Coordinated shape changes such as reciprocal deformation in synergists or directional bulging in pennate muscles may represent strategies to maximise spatial efficiency and protect anatomical structures. These spatial-functional adaptations represent an elegant solution to spatial constraints within functional systems and may have implications for both biological understanding and translation applications.
7. A better understanding of variable gearing is particularly important in clinical contexts. Many conditions involving disuse, whether due to immobilisation, injury, ageing, or neurological or musculoskeletal experiments, lead to substantial alterations in muscle architecture and tissue properties and are often accompanied by rapid declines in muscle power and functional capacity. Other physiological states may conversely enhance muscle performance via changes in tissue hydration or stiffness. Evidence suggests that ageing and disuse can attenuate or abolish fascicle reorientation and impair variable gearing, while acute electrically induced maximal contraction appears to enhance performance and increase the magnitude of fascicle reorientation. As such, influences on variable gearing may help explain these changes and offer a framework for interpreting muscle function loss across a range of conditions, including ageing, disease, and physical training or detraining. Advancing our understanding of these phenomena may provide a basis for understanding human muscle function and movement performance, and reveal novel targets for future intervention through exercise, nutrition, or pharmacological strategies.

## X. Acknowledgements

MDP was supported by funding from the Australasian Society of Human Biology, the Comparative Neuromuscular Biomechanics Group, and a PhD scholarship from the Australian Government Research Training Program. JAA acknowledges financial support from Simon Fraser University, Canada, through the Graduate Dean’s Entrance Scholarship. JMW was supported by a Discovery Grant from the Natural Sciences and Engineering Research Council (NSERC) of Canada.

